# A drug-screening platform based on organotypic cultures identifies vulnerabilities to prevent local relapse and treat established brain metastasis

**DOI:** 10.1101/2020.10.16.329243

**Authors:** Lucía Zhu, Natalia Yebra, Diana Retana, Lauritz Miarka, Elena Hernández-Encinas, Carmen Blanco-Aparicio, Sonia Martínez, Riccardo Soffietti, Luca Bertero, Paola Cassoni, Tobias Weiss, Javier Muñoz, Juan Manuel Sepúlveda, Ángel Pérez-Núñez, Aurelio Hernández-Laín, Yolanda Ruano, Oscar Toldos, Eduardo Caleiras, Diego Megías, Osvaldo Graña-Castro, Carolina Nör, Michael D. Taylor, Lorena Cussó, Manuel Desco, Michael Weller, Joaquín Pastor, Manuel Valiente

**Affiliations:** Brain Metastasis Group; CNIO; Madrid, 28003; Spain; Experimental Therapeutics Programme; CNIO; Madrid, 28003; Spain; Department of Neuro-Oncology; University and City of Health and Science Hospital; Turin, 10126; Italy; Department of Medical Sciences/ Pathology Unit; University of Turin; Turin, 10124; Italy; Department of Neurology and Brain Tumor Center; University Hospital Zurich and University of Zurich; Zurich, 8091; Switzerland; Proteomics Unit; CNIO; Madrid, 28003; Spain. ProteoRedISCIII; Neuro-Oncology Unit; Hospital Universitario 12 de Octubre; Madrid, 28041; Spain; Neurosurgery Unit; Hospital Universitario 12 de Octubre; Madrid, 28041; Spain; Neuropathology Unit; Hospital Universitario 12 de Octubre; Madrid, 28041; Spain; Histopathology Unit; CNIO; Madrid, 28003; Spain; Confocal Microscopy Unit; CNIO; Madrid, 28003; Spain; Bioinformatics Unit; CNIO; Madrid, 28003; Spain; Developmental and Stem Cell Biology Program and The Arthur and Sonia Labatt Brain Tumour Research Centre; The Hospital for Sick Children; Toronto, M5G 1X8; Canada; Departamento de Bioingeniería e Ingeniería Aeroespacial; Universidad Carlos III de Madrid; Madrid, 28911; Spain; Instituto de Investigación Sanitaria Gregorio Marañón; Madrid, 28007; Spain; Centro de Investigación Biomédica en Red de Salud Mental (CIBERSAM); Spain; Unidad de Imagen Avanzada; Centro Nacional de Investigaciones Cardiovasculares (CNIC); Madrid, 28003; Spain

**Keywords:** Metastasis, drug-screen, organotypic cultures, brain metastasis, resistance, biomarker, proteomics, patient-derived, local relapse, drug-repurposing

## Abstract

Exclusion of brain metastases from clinical trials is a major cause of the limited therapeutic options for this growing population of cancer patients. Here, we report a medium-throughput drug-screening platform (METPlatform) based on organotypic cultures that allows to evaluate inhibitors against metastases growing *in situ*. By applying this approach to brain metastasis, we identified several hits from a library of FDA approved inhibitors and others being tested in clinical trials. A blood-brain barrier permeable HSP90 inhibitor showed high potency against mouse and human brain metastases at clinically relevant stages of the disease, including a novel model of local relapse after neurosurgery. Furthermore, *in situ* proteomic analysis applied to organotypic cultures with metastases treated with the chaperone inhibitor revealed novel biomarkers in human brain metastasis and actionable mechanisms of resistance. Our work validates METPlatform as a potent resource for metastasis research integrating drug-screening and unbiased omic approaches that is fully compatible with human samples. We envision that METPlatform could be established as a clinically relevant strategy to personalize the management of metastatic disease in the brain and elsewhere.

**Summary:** Systemic spread of cancer continues to be the key aspect associated with lethality. In this publication, Zhu et al. describes a drug-screening platform specifically designed to study vulnerabilities of metastasis when colonizing secondary organs and demonstrates its value in difficult-to-treat brain metastasis using new models and patient-derived samples.

## Introduction

Incidence of brain metastasis continue to increase and yet, current therapies available for patients with disseminated cancer cells in their central nervous system (CNS) yield limited efficacy and fail to improve survival (Moravan et al., 2020; Suh et al., 2020; Valiente et al., 2018).

Existing evidence shows that these patients would qualify as those without brain metastasis to be included in clinical trials with systemic anti-cancer drugs (Tsimberidou et al., 2011). In addition, during the past years, there have been recurrent efforts to improve clinical trial design and management specifically concerning this patient population (Lin et al., 2013a, 2013b, 2015). However, complete exclusion of patients with active CNS disease remains an unsolved issue (Arvold et al., 2016). Retrospective analysis of clinical trials evaluating CNS efficacy of systemic agents generally overestimate the impact on brain metastases due to trial design limitations. For instance, the poor dissociation between the efficacy derived from the drug under study and the benefits provided by previously administered therapies (i.e. radiotherapy) is a good example. In addition, a biased assessment towards stable over progressive disease due to the lack of more sensitive criteria to evaluate the local impact of a given therapy reflect some of these limitations (Camidge et al., 2018). As a result of all previous considerations, information regarding CNS clinical efficacy of most anti-cancer agents that are FDA-approved or in clinical trials is limited. Thus, the use of preclinical models to explore therapeutic vulnerabilities and the subsequent analysis of CNS activity of adequate pharmacological agents are crucial to promote urgently needed prospective clinical trials that include patients with brain metastases (Camidge et al., 2018).

Drug-screening using mouse models that faithfully recapitulate the clinical phenotype imposes high demand of economic costs and resources (Gao et al., 2015) that are unaffordable by most academical research institutions. On the other hand, cell-based assays lack the contribution of the tumor-associated microenvironment, which has gained relevance in the context of response to therapy during recent years (Hirata and Sahai, 2017). Indeed, the brain microenvironment is a key aspect in the biology of CNS metastasis (Boire et al., 2020) that has been demonstrated to limit therapeutic benefits of systemic therapy (Chen et al., 2016).

To overcome limitations of both *in vivo* and *in vitro* approaches, we report an organotypic culture-based drug-screening system: METPlatform. We use this strategy to evaluate the impact of different therapeutic agents on brain metastases *in situ*, thus identifying biologically relevant drug candidates in a rapid and cost-effective manner.

Brain organotypic cultures have been used in cancer research due to their ability to mimic the progression of metastatic disease in the brain (Zhu and Valiente). They resemble both early (Er et al., 2018; Valiente et al., 2014) and advanced clinically relevant stages of the disease (Priego et al., 2018). Their versatility allows exploring diverse functional and mechanistic insights of brain metastasis, including the interaction between cancer cells and different components of the microenvironment using genetic or pharmacologic approaches (Er et al., 2018; Priego et al., 2018; Valiente et al., 2014). However, to the best of our knowledge, their use for drug-screening has not been reported. We describe here the use of brain organotypic cultures for performing a medium-throughput screening using an in-house library of anti-cancer agents, FDA-approved or under clinical development (Bejarano et al., 2019), with unknown or limited information regarding their activity in the CNS. In addition to other hits, METPlatform identified inhibitors of heat shock protein 90 (HSP90) as a potential therapy for brain metastasis.

HSP90 is a molecular chaperone required for correct protein folding, intracellular disposition and proteolytic turnover of its client proteins, and therefore essential for cellular proteostasis (Schopf et al., 2017). It is heavily exploited by cancer cells not only to maintain numerous pro-survival oncoproteins and transcription factors, but also to buffer proteotoxic stress induced during oncogenic transformation and progression (Whitesell and Lindquist, 2005) as well as to regulate mechanisms of immune evasion (Fionda et al., 2009; Kawabe et al., 2009). High HSP90 expression levels have been correlated with poor prognosis in all subtypes of breast cancer patients (Dimas et al., 2018; Pick et al., 2007), several independent cohorts of non-small cell lung cancer (NSCLC) patients (Gallegos Ruiz et al., 2008), and in colorectal cancer (Kim et al., 2019).

Here we report the potent anti-metastatic activity of a second generation HSP90 inhibitor, DEBIO-0932, in clinically relevant stages of systemic disease affecting the brain both in experimental and human metastases. In addition, we use METPlatform to study the underlying biology downstream of HSP90 inhibition using unbiased proteomics, which has allowed us to identify potential mediators of brain metastasis and effective combination strategies to overcome resistance.

## Results

### A chemical library applied to METPlatform identifies potential vulnerabilities of brain metastasis

Given our interest in targeting clinically relevant stages of brain metastasis, we used METPlatform to study vulnerabilities of macrometastases. The human lung adenocarcinoma brain metastatic (BrM) cell line H2030-BrM (Nguyen et al., 2009) was injected intracardially into athymic nude mice to obtain fully established brain macrometastases at clinical endpoint of the animals. Brains were processed into organotypic cultures and the efficacy of the anti-tumoral library (Table 1) was evaluated at a concentration of 10 μM (Fig. 1 A). Given the expression of luciferase and GFP in the H2030-BrM model (Nguyen et al., 2009), the impact of specific inhibitors on the viability of brain metastases in organotypic cultures were assessed by bioluminescence imaging (BLI) and immunofluorescence against GFP in comparison to DMSO treated cultures. We used a PI3K inhibitor, BKM120, as an internal positive control in our experiments due to the known involvement of this signaling pathway and therapeutic benefit in brain metastasis (Brastianos et al., 2015; Nanni et al., 2012; Pistilli et al., 2018). The analysis of the drug-screen provided us with 17 hits: carfilzomib (ref#1), dovitinib (ref#9), trametinib (ref#22), mitomycin C (ref#39), GSK2126458 (ref#44), AT7519 (ref#52), CNIO-DUAL (ref#56), sorafenib (ref#59), geldanamycin (ref#60), SN-38 (ref#72), bortezomib (ref#84), KU-57788 (ref#87), CNIO-TRIPLE (ref#104), crizotinib (ref#106), CNIO-ATR (ref#107), pazopanib (ref#110), linifanib (ref#113) out of 114 compounds tested (Fig. 1, B and C, Table 1). Hits were defined as compounds able to reduce in 80% or more the bioluminescence values that correspond to controls treated with DMSO. This threshold was confirmed to be a good correlate of compromised viability based on a complementary histological analysis (Fig. 1 C). Notice that, in spite of reproducing the efficacy of BKM120, METPlatform identified hits that are superior to an established inhibitor against brain metastasis.

**Figure 1.**
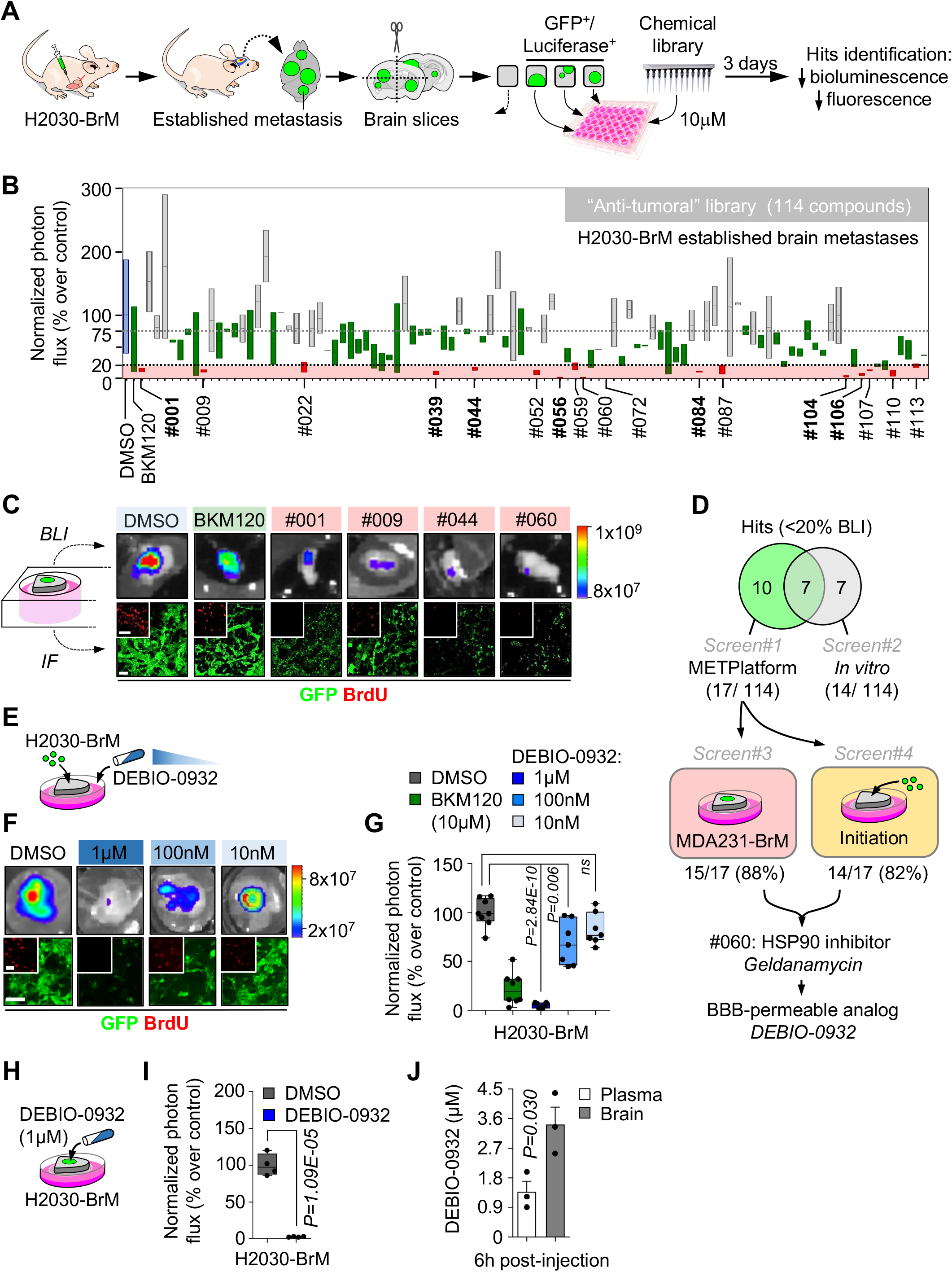
A chemical library applied to METPlatform identifies potential vulnerabilities of brain metastasis. **(A)** Schema of the experimental design. **(B)** Quantification of the bioluminescence signal emitted by established H2030-BrM brain metastases in each organotypic culture at day 3 normalized by their initial value at day 0 (before the addition of DMSO or any compound). The final value in the graph is normalized to the organotypic cultures treated with DMSO. Blue: DMSO-treated organotypic cultures; red: hits, compounds with normalized BLI ≤20%; green: BKM120 and compounds with similar efficacy to BKM120; gray: compounds that do not reduce BLI values. Values are shown in box-and-whisker plots where the line in the box corresponds to the mean. Whiskers go from the minimum to the maximum value (n=28 DMSO; n=21 BKM120-treated organotypic cultures; each experimental compound of the library was assayed by duplicate, 8 independent experiments). Hits highlighted in bold are common to those obtained in the *in vitro* screening (Fig. S1 A). **(C)** Representative images of bioluminescence (BLI) and histology of organotypic cultures with established brain metastases from H2030-BrM treated with DMSO, BKM120 or the indicated hits. Cancer cells are in green (GFP) and proliferative cells are in red (BrdU). Scale bar, 75 μm. **(D)** Venn diagram showing the number of hits *ex vivo* (17) and *in vitro* (14) and common to both approaches (7). Compounds tested in additional screens (screen#3: established MDA231-BrM breast cancer brain metastasis; and screen#4: metastasis initiation H2030-BrM) only include those considered as hits *ex vivo* in panel (B). Number of hits in each screen are indicated over the total number of hits obtained in screen#1 (B). **(E)** Schema of the experimental design. Organotypic cultures with H2030-BrM cells mimicking the early steps of colonization were used to perform dose-response optimization with DEBIO-0932. **(F)** Representative BLI and histology of organotypic cultures with H2030-BrM cancer cells treated with DMSO or decreasing concentrations of DEBIO-0932. Scale bar, 100 μm; high magnification: 50 μm. **(G)** Quantification of the bioluminescence signal emitted by each condition shown in (F) at day 3 normalized by the initial value obtained at day 0 and normalized to the organotypic cultures treated with DMSO. Day 0 is considered 12-16h after the addition of cancer cells and treatment or DMSO. Values are shown in box-and-whisker plots where each dot is an organotypic culture and the line in the box corresponds to the median. Whiskers go from the minimum to the maximum value (n=8 DMSO, n=8 BKM120 and n=7 per concentration of DEBIO-0932-treated organotypic cultures, 2 independent experiments). *P* value was calculated using two-tailed t-test. **(H)** Schema of the experimental design. Organotypic cultures with H2030-BrM established metastases were used to test the efficacy of DEBIO-0932. **(I)** Quantification of the bioluminescence signal emitted by H2030-BrM established metastases in organotypic cultures at day 3 normalized by the initial value obtained at day 0 and normalized to the organotypic cultures treated with DMSO. Day 0 is considered right before addition of the treatment or DMSO. Values are shown in box-and-whisker plots where each dot is an organotypic culture and the line in the box corresponds to the median. Whiskers go from the minimum to the maximum value (n=4 organotypic cultures per experimental condition, 2 independent experiments). *P* value was calculated using two-tailed t-test. **(J)** Quantification of the concentration of DEBIO-0932 reached in animals harboring H2030-BrM established brain metastases 6h after oral administration of DEBIO-0932 at 160 mg/kg. The concentration was measured in both the plasma and the brain for each mouse. Values are shown as mean + s.e.m. (n=3 mice per experimental condition). *P* value was calculated using two-tailed t-test.

**Table 1.**
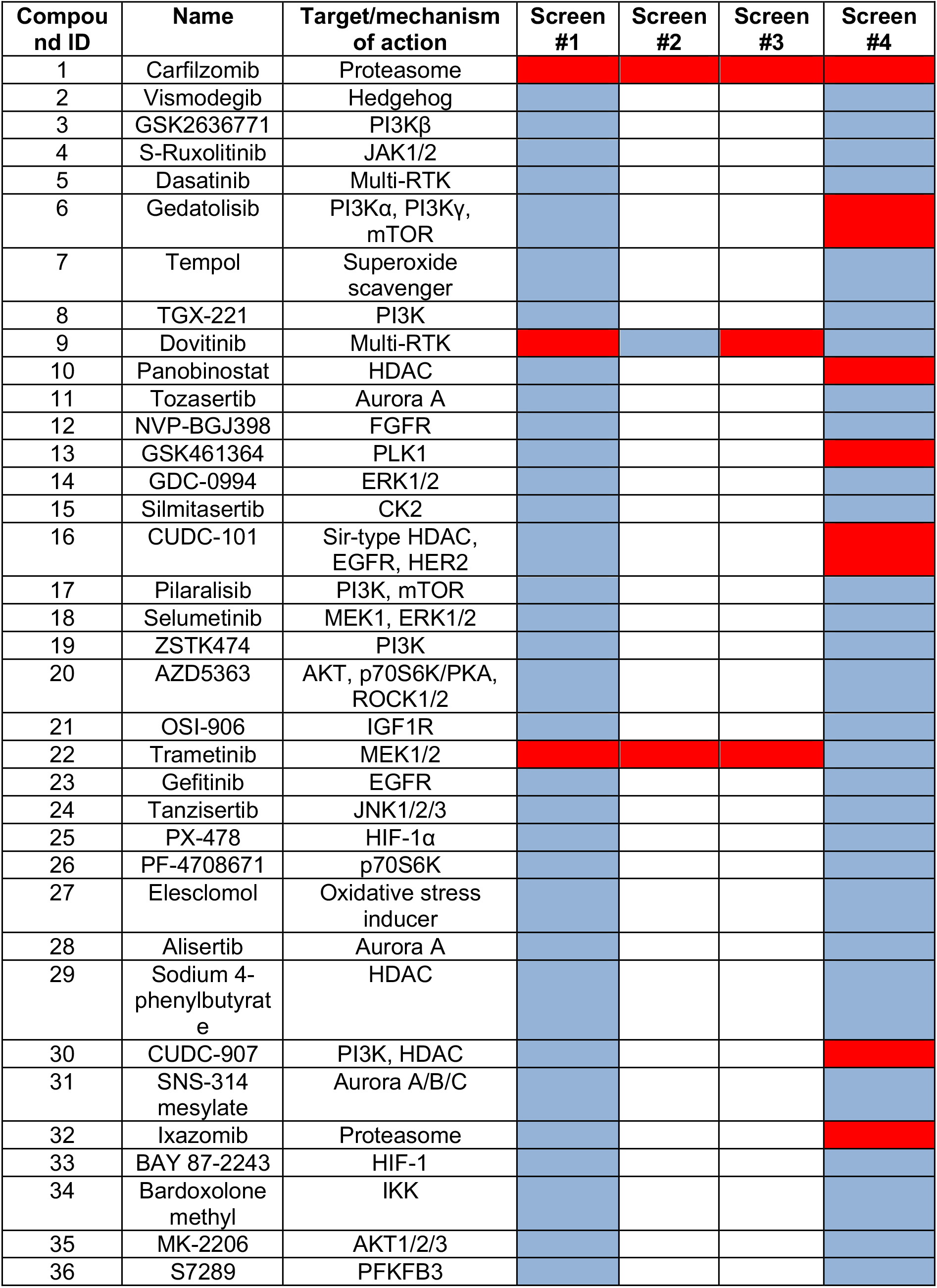
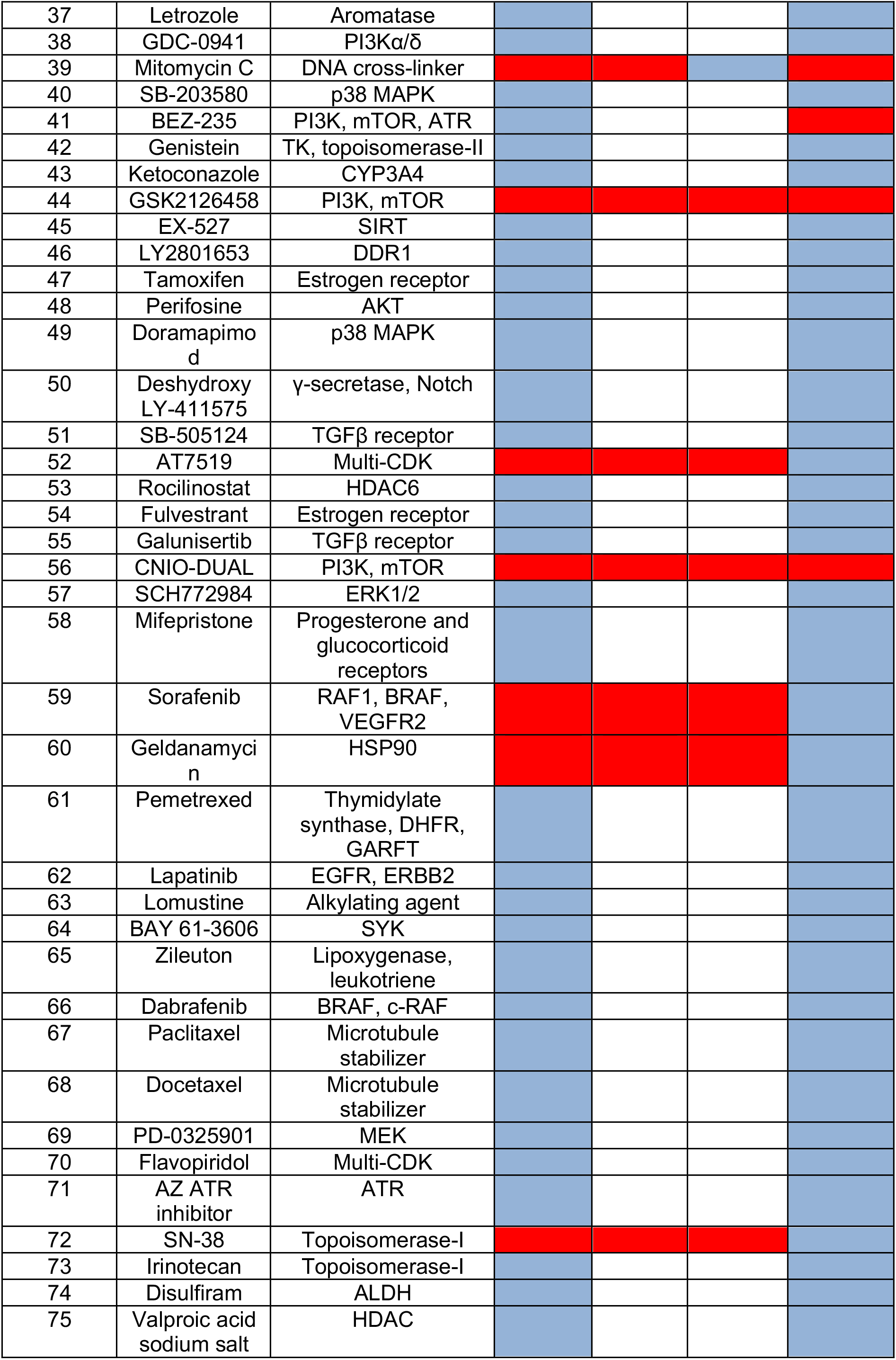
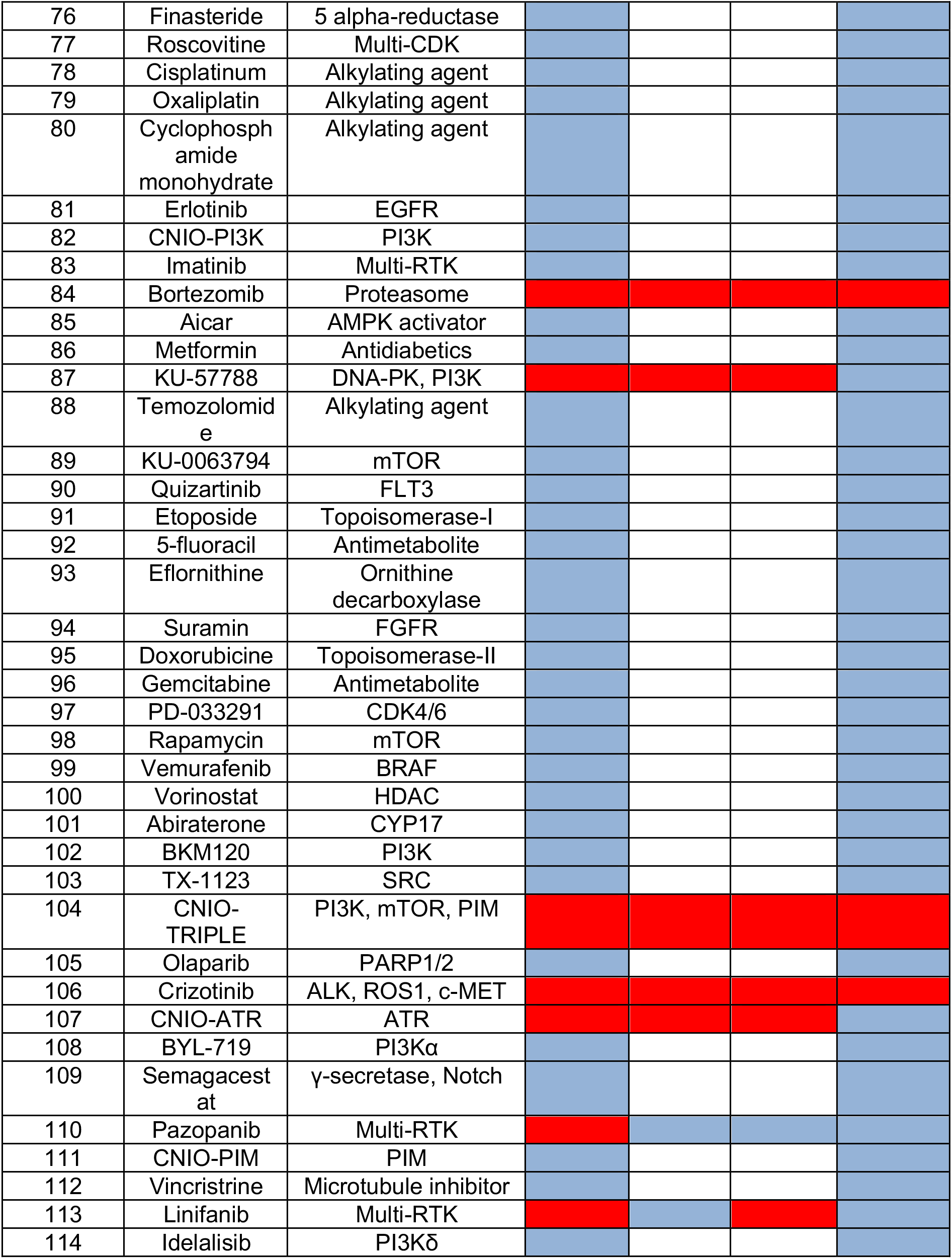
Antitumoral library and drug-screens. 114 compounds present in this library have been applied to different screens. In screen#1 we have tested 114 compounds in METPlatform using lung adenocarcinoma (H2030-BrM) established metastases. In screen#2 we have tested hits obtained from screen#1 in METPlatform using lung adenocarcinoma (H2030-BrM) brain metastases representing the initial stages of organ colonization. In screen#3 we have tested hits obtained from screen#1 in METPlatform using breast adenocarcinoma/ triple negative subtype (MDA231-BrM) established brain metastases. Screen#4 corresponds to the use of 114 compounds on H2030-BrM cells *in vitro*, which represents traditional strategies to compare with METPlatform. Red: labels a hit in a specific screen. Blue: labels a tested compound that has not been considered as a hit in a given screen. White: compounds not tested in a given screen.

To compare METPlatform with a traditional cell-based assay as a drug-screening platform, we applied the same chemical library to H2030-BrM cells cultured *in vitro* (Fig. S1 A). Interestingly, after applying the same criteria based on luminescence, only 7 out of 14 hits obtained *in vitro* were part of the 17 hits obtained with METPlatform (Fig. 1 D, Fig. S1 C, Table 1). Thus, METPlatform selected hits that would not have been considered as such in an *in vitro* approach.

We extended our *ex vivo* drug-screen to a triple negative breast cancer brain metastasis model, MDA231-BrM (Bos et al., 2009), to identify vulnerabilities regardless the origin of the primary tumor. Out of the 17 hits tested, 15 of them decreased the viability of cancer cells in 80% or more as measured by BLI (Fig. 1 D, Fig. S1 B, Table 1). In addition, we used METPlatform to analyze whether any hit also scored not only against advanced stages of the disease when metastases are fully established (Fig. 1 B), but also against the initial steps of organ colonization, which could be mimicked *ex vivo* by plating cancer cells on top of tumor-free organotypic brain cultures (Valiente et al., 2014). Interestingly, 14 out of 17 hits inhibited both early and advanced stages of brain metastasis (Fig. 1 D, Table 1), which suggests that these compounds may not only be effective treating, but also preventing metastasis outgrowth by acting on the initiation of organ colonization. On the other hand, reported differences in the biology of initial and established brain metastases (Priego et al., 2018; Valiente et al., 2014) could be exploited therapeutically by interrogating those hits only scoring in one or another stage of colonization (dovitinib (ref#9), pazopanib (ref#110) and linifanib (ref#113)) (Table 1).

Finally, METPlatform also allows simultaneous evaluation of the potential toxicity derived from selected compounds on non-cancer cell types and in different organs. For instance, the use of specific markers for various cell types present in the brain, such as neurons and endothelial cells, allowed us to discard a major unselective cytotoxicity in this organ (Fig. S1 D). In contrast, evaluation of reported sensitive organs confirmed the ability of the drug-screening platform to reproduce clinical toxicity (i.e. hepatotoxicity) (Fig. S1 E) (Supko et al., 1995).

Altogether, our results support METPlatform as a comprehensive and more informative drug-screening platform in the context of metastasis compared to conventional cell-based assays (Fig. 1 D, Table 1).

In order to select compounds for further validation we focused on those targeting not only established metastasis from different cancer types but also initial stages of organ colonization (Fig. 1 D, Fig. S1 B, Table 1). Out of this selection we then focused on those that, although with significant inhibitory activity *in vitro* (Fig. S1 F), did not score as hits in this condition (Fig. S1, A and C). With this selection criterium we wanted to evaluate the potential of METPlatform to select hits working *in vivo*. Seven hits fulfilled the criteria: trametinib (ref#22), AT7519 (ref#52), sorafenib (ref#59), geldanamycin (ref#60), SN-38 (ref#72), KU-57788 (ref#87), CNIO-ATR (ref#107). Unfortunately, METPlatform has no capacity to score blood-brain barrier (BBB) permeability and indeed we failed to recognize this property among these compounds, suggesting that, when METPlatform is applied to metastasis in the brain, a previous step to prioritize BBB-permeable compounds should be incorporated to design the compound library (Saxena et al., 2019). Given the improved efficacy of brain permeable compounds to target metastasis in this organ (Osswald et al., 2016), we looked for alternative inhibitors focused on the targets identified. DEBIO-0932, a second generation HSP90 inhibitor, has an improved toxicity profile in comparison with geldanamycin, increased bioavailability and, more importantly, a remarkable ability to cross the BBB (Bao et al., 2009; Supko et al., 1995). As geldanamycin, DEBIO-0932 blunted the viability of initial and established brain metastases from lung (H2030-BrM) and breast (MDA231-BrM) cancer models (Fig. 1, E-I, Fig. S1 G) in *ex vivo* assays. Furthermore, the concentration reached by DEBIO-0932 in a brain affected by metastases (Fig. 1 J) is above the therapeutic levels as determined *ex vivo* (Fig. 1, E-I).

Altogether, METPlatform identified potential vulnerabilities of brain metastasis that could be targeted with inhibitors that are currently available or under clinical trials. Among them we identified DEBIO-0932 as a potent inhibitor of brain metastases viability *ex vivo* that is able to accumulate in the brain at therapeutic concentrations.

### Brain metastases are positive for HSP90

Before testing the potential benefits of DEBIO-0932 *in vivo*, we evaluated the presence of its target in experimental and human brain metastasis. To evaluate HSP90 levels *in situ*, we performed tissue immunofluorescence in four experimental brain metastasis models from both human and mouse origin, characterized by different oncogenic drivers and derived from breast cancer, lung cancer and melanoma, which are the most frequent sources of brain metastasis (Valiente et al., 2020). Established brain macrometastases obtained 5 weeks (human cancer cell lines) or 2 weeks (mouse cancer cell line, B16/F10-BrM) post-injection (Valiente et al., 2020) showed high HSP90 levels in cancer cells. In sharp contrast, the metastasis-associated microenvironment has some positive cells but with lower intensity than metastases (Fig. 2, A and B). Finally, only discrete areas (hippocampus, isocortex) of normal brains have some positivity (Fig. 2, A and B).

**Figure 2.**
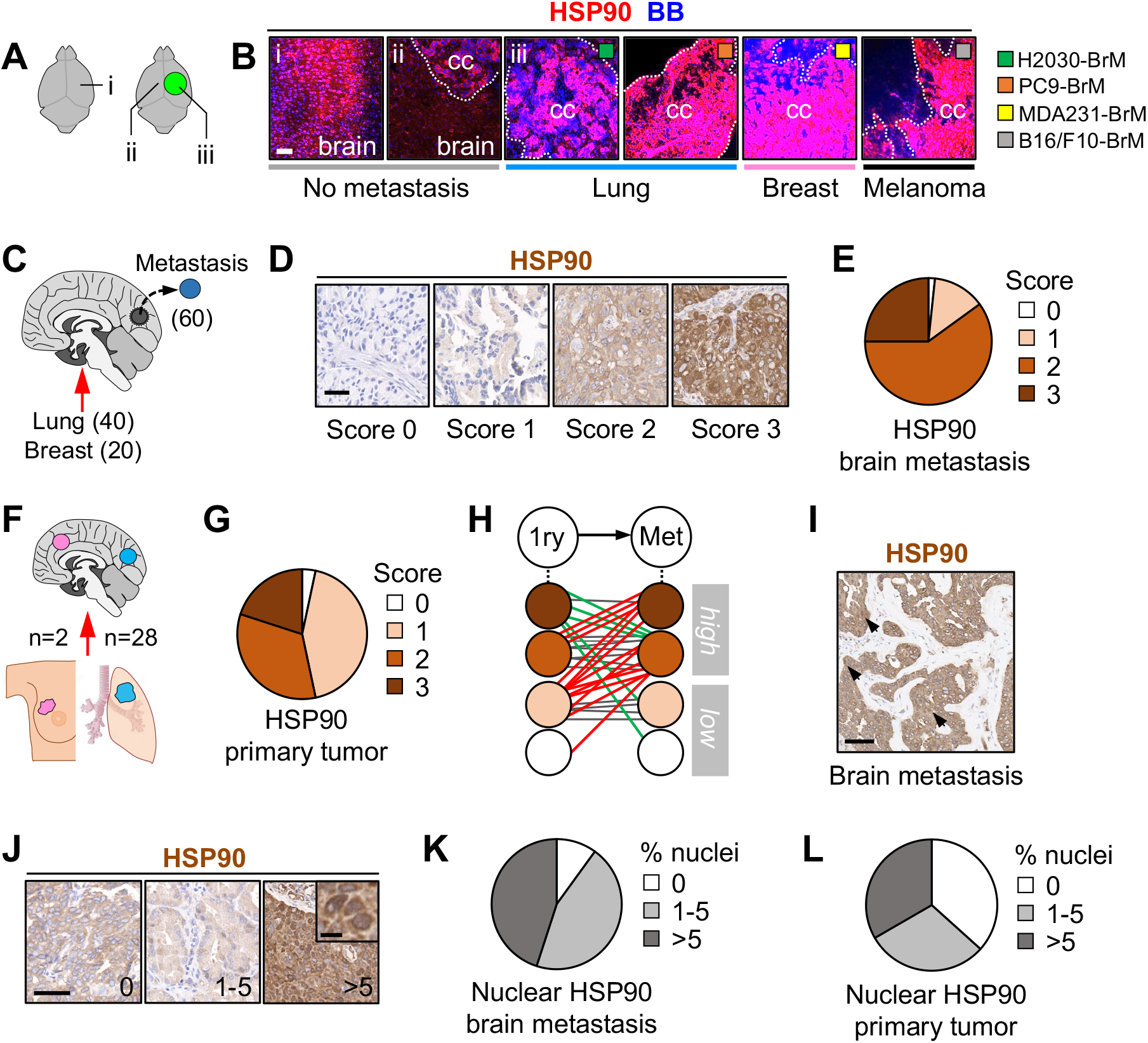
Brain metastases are positive for HSP90. **(A-B)** Immunofluorescence against HSP90 in mouse brains without (i) or with established metastases (ii-iii) from different primary origins and oncogenomic profiles. Dotted lines delineate the metastasis (cc: cancer cells). BB: bisbenzamide. Scale bar, 75 μm. **(C)** Immunohistochemistry against HSP90 was performed in human brain metastases (n=60) from lung (40 cases) and breast cancer (20 cases). **(D)** Representative human brain metastases showing different intensities or scores for HSP90. Scale bar, 50 μm. **(E)** Quantification of HSP90 in human brain metastases. 59 out of 60 (98%) showed positive staining of HSP90 in the tumor, 15 (25%) scored with 3 (strong), 36 (60%) with 2 (moderate) and 8 (13%) with 1 (weak) according to the signal intensity of HSP90 in the cytoplasm of cancer cells. **(F)** Human brain metastases (n=30) and their matched primary tumors (n=28 lung and n=2 breast) were evaluated and compared for HSP90 expression by immunohistochemistry. **(G)** Quantification of HSP90 in human primary tumors. 29 out of 30 (97%) showed positive staining of HSP90 in the tumor, 6 (20%) scored with 3 (strong), 10 (34%) with 2 (moderate) and 13 (43%) with 1 (weak) according to the signal intensity of HSP90 in the cytoplasm of cancer cells. **(H)** Schema showing HSP90 scores in matched pairs of primary tumor and brain metastasis. Red: increase of HSP90 score from primary to brain metastasis; green: decrease of HSP90 score; gray: no changes in HSP90 score. **(I-J)** Representative human brain metastases showing different percentages of nuclear HSP90. Scale bar, 50 μm; low magnification, 100 μm; high magnification, 10 μm. **(K)** Quantification of nuclear HSP90 in human brain metastases. 54 out of 60 (90%) showed positive nuclear HSP90 in the tumor. 27 (45%) showed 1-5% (moderate) and 27 (45%) showed >5% (high) of nuclear HSP90. **(L)** Quantification of nuclear HSP90 in human primary tumors. 19 out of 30 (63%) showed positive nuclear HSP90 in the tumor. 9 (30%) showed 1-5% (moderate) and 10 (33%) showed >5% (high) of nuclear HSP90.

Additionally, 60 paraffin-embedded human brain metastases from NSCLC (40 samples) and breast adenocarcinoma (20 samples) were stained with anti-HSP90 by immunohistochemistry and blindly evaluated and scored by a pathologist (Fig. 2 C). 98% of brain metastases were positive for HSP90, with 85% of them showing moderate or strong staining of the protein (score ≥2, HSP90^high^) (Fig. 2, D and E), which is a value higher than previous reports on primary tumors (Gallegos Ruiz et al., 2008; Kim et al., 2019; Pick et al., 2007). To investigate this possibility, we scored 30 matched primary tumors (Fig. 2 F) and confirmed a lower percentage (54%) of samples scoring as HSP90^high^ in comparison to brain metastases (Fig. 2 G). When comparing matched pairs of a primary tumor and a brain metastasis, 13/30 (43%) brain metastases had increased HSP90 levels compared to the primary tumor, from which 10/13 (77%) switched from HSP90^low^ (score ≤1) to HSP90^high^ (score ≥2). 12/30 (40%) matched pairs showed equal HSP90 levels, however, 8/12 (67%) cases were HSP90^high^ in the primary tumor to start with. Out of the 5/30 (17%) brain metastases with lower HSP90 than the corresponding primary tumor, 3/5 (60%) cases still remained within the HSP90^high^ category and only 2/5 (40%) switched from HSP90^hi^g^h^ to HSP90^low^ (Fig. 2 H).

Although HSP90 is primarily a cytoplasmic protein, several studies have described its role in nuclear events such as transcriptional processes, chromatin remodeling and DNA damage (Antonova et al., 2019; Trepel et al., 2010). Moreover, increased nuclear HSP90 correlated positively with poor survival and distant metastasis in NSCLC patients (Su et al., 2016). Interestingly, we found nuclear staining of HSP90 in 90% of brain metastasis samples (Fig. 2, I-K), with 45% of them scoring as HSP90^high^ (>5% of positive nuclei out of total tumor) according to a previously described criteria (Su et al., 2016) (Fig. 2 K). Similar to the previous analysis, we found fewer primary tumors (63%) positive for nuclear HSP90, with 33% of them scoring HSP90^high^ (Fig. 2 L). Nevertheless, due to the prevalent low percentage of positive nuclei observed in most samples (Fig. 2 I), we were not able to accurately assess a potential enrichment of nuclear HSP90 in brain metastases compared to their paired primary tumor.

Taken together, our results demonstrate that high levels of HSP90 is a frequent finding among human brain metastasis independently of the primary tumor. Indeed, a clear tendency to maintain or further increase the levels of this protein is evident when compared to matched primary tumors. Overall, these results support potential functional implications of HSP90 in human brain metastasis.

### Inhibition of HSP90 impairs clinically-relevant stages of brain metastasis

We used DEBIO-0932 in established and novel preclinical models to study whether the results obtained with METPlatform could be translated *in vivo*.

Brain metastases were induced by intracardiac inoculation of H2030-BrM cells (Nguyen et al., 2009). Two weeks after injection, we confirmed the presence of established metastases in the brain using BLI, histology and magnetic resonance imaging (MRI), which is used to diagnose brain tumors in patients (Fig. 3 A). DEBIO-0932 administration at 160mg/kg during the following 3 weeks significantly impaired the growth of both brain metastases and extracranial lesions as measured by BLI (Fig. 3, B-D) by targeting HSP90 (Fig. S3, A-D). These results were confirmed by brain and thorax *ex vivo* BLI (Fig. 3, E-G) as well as histological quantification of dissected brains (Fig. S3, E and F) at the endpoint of the experiment, 5 weeks after cancer cell inoculation. Of note, we did not observe significant weight loss in treated animals compared to the control group (Fig. S3 G), discarding major toxicities of DEBIO-0932 under the therapeutic regimen followed.

**Figure 3.**
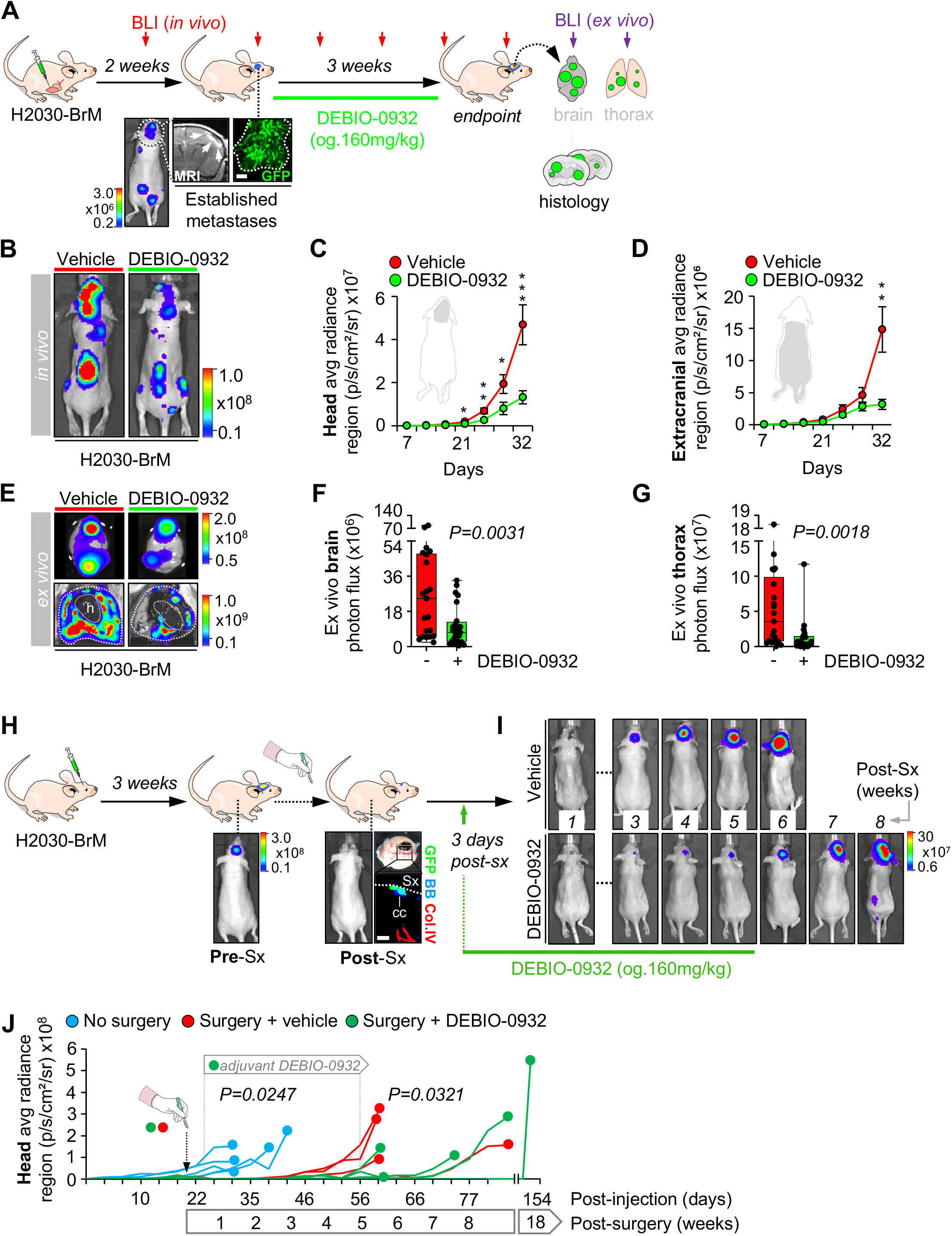
Inhibition of HSP90 impairs clinically-relevant stages of brain metastasis. **(A)** Schema of the experimental design. H2030-BrM cells were inoculated intracardially into nude mice and established brain metastases were detected 2 weeks after by BLI, MRI (arrows) and histology (GFP+ cancer cells). DEBIO-0932 was administered orally at 160 mg/kg for 3 weeks and *ex vivo* BLI of brains and thoracic regions were analyzed. Brains were processed for histological analysis. Scale bar, 100 μm. **(B)** Representative images of vehicle and DEBIO-0932 treated mice 5 weeks (experimental endpoint) after intracardiac inoculation of H2030-BrM cells. **(C-D)** Quantification of metastatic progression as measured by *in vivo* BLI of head (C) and extracranial region (D) of animals. Values are shown as mean ± s.e.m. (n=23 vehicle and n=25 DEBIO-0932 treated mice, 3 independent experiments). *P* value was calculated using two-tailed t-test (*P* values: \**P*<0.05, \*\**P*<0.01, \*\*\**P*<0.001). **(E)** Representative images of brains and thorax from vehicle and DEBIO-0932 treated mice at the endpoint of the experiment. **(F-G)** Quantification of *ex vivo* BLI of brains (F) and thoracic regions (G) at the endpoint of the experiment. Values are shown in box-and-whisker plots where every dot represents a different animal and the line in the box corresponds to the median. Whiskers go from the minimum to the maximum value. (n=21 vehicle and n=24 DEBIO-0932 treated mice, 3 independent experiments). *P* value was calculated using two-tailed t-test. **(H)** Schema of experimental design. H2030-BrM cells were implanted intracranially into nude mice and established brain metastases were surgically resected. Remaining cancer cells (GFP+) were found in the surgical bed, in which endothelial cells are present (Col.IV+). DEBIO-0932 was administered orally at 160 mg/kg 3 days later and during 5-6 weeks. Scale bar, 25 μm. Sx: surgery. BB: bisbenzamide. **(I)** Representative images of vehicle and DEBIO-0932 treated mice after neurosurgery until experimental endpoint at 6 and 8 weeks for vehicle and DEBIO-0932 treated mice, respectively. **(J)** Quantification of brain tumor progression as measured by *in vivo* BLI of head region in animals without surgery, with surgery and vehicle or DEBIO-0932. DEBIO-0932 treatment was initiated 3 days after surgery, which was applied 3 weeks postinjection of BrM cells, and maintained for 5-6 weeks after local treatment. Each line represents an animal (n=5 without surgery, n=4 surgery + vehicle and n=5 surgery + DEBIO-0932 treated mice). *P* value was calculated using two-tailed t-test (No surgery *versus* surgery + vehicle (day 32), *P*=0.0247; surgery + vehicle *versus* surgery + DEBIO-0932 (day 53), *P*=0.0321).

Approximately 20-40% of patients with brain metastasis receive neurosurgery. However, local relapse is a common occurrence that limits the benefits of an otherwise successful local therapy (Nahed et al., 2019). To investigate whether DEBIO-0932 is able to prevent this clinically relevant situation for which there is no established standard of care, we developed a first-in-class preclinical model of local relapse after brain metastasis neurosurgery.

We modelled single brain macrometastasis by intracranial implantation of H2030-BrM cells. This strategy facilitates the surgical approach avoiding non-operable brains with multiple secondary tumors or surgically non-accessible locations of metastasis (Valiente et al., 2020). Microsurgical resection of the metastasis guided by GFP was performed when BLI values reached 10^7^ photons/sec/cm^2^/steradian (Fig. 3, H and J, Fig. S3, H and I). Successful resection of the bulk tumor was confirmed in real-time by the absence of macroscopically detectable GFP+ cancer cells and almost complete post-surgical reduction of BLI *in vivo* (Fig. 3, H and J, Fig. S3 I). However, the presence of single cancer cells left behind was also evident by microscopic analysis of the borders of the surgical bed one day after completing the local treatment (Fig. 3 H). These cancer cells are presumably responsible of the local relapse that affected all treated mice since tumors always reappeared within the same area where mass debulking was initially applied (Fig. S3 K). Full development of relapsed tumors occurred 5 to 6 weeks after surgery (Fig. 3, I and J), which double the survival time compared to those animals without the local treatment (Fig. S3 J). We validated that HSP90 coding genes and members of the heat shock response pathway were unaltered in relapsed tumors (Fig. S3 L). However, differences between resected and relapsed tumors do exist. Gene Set Enrichment Analysis of the transcriptomes from relapsed versus matched resected tumors showed downregulated signatures related to cell cycle and proliferation and enrichment in those related to vascular co-option, a key mechanism during the early stages of organ colonization (Er et al., 2018; Valiente et al., 2014), and cytokine and integrin signaling (Fig S3 M, Table 2).

**Table 2.** Top 25 upregulated and downregulated signatures in relapsed versus resected brain metastasis transcriptome. GSEA analysis has been applied to the transcriptome of matched resected and relapsed metastases. Pink: signatures belonging to cell cycle, proliferation. Light blue: signatures belonging to DNA, RNA or protein processes. Yellow: signatures belonging to vascular co-option (*). Dark blue: signatures belonging to cytokine signaling. Light green: signatures belonging to integrin signaling.

| UP |  |  |  |  |  |
| --- | --- | --- | --- | --- | --- |
| ID path way | Dataset | Pathway | NES | Nominal p-value | FDR q-value |
| #1 |  | CO-OPTION<br>UP_GENES_LOG2FC_1 | 3.6213484 | 0 | 0 |
| #2 | OncogenicSignatures.GseaPreranked.1585307079701 | STK33_NOMO_UP | 2.6241393 | 0 | 0.0035213 |
| #3 | CanonicalPathways.GseaPreranked.1585307356087 | REACTOME_CYTOKINE_SIGNALING_IN_IMMUNE_SYSTEM | 2.4835162 | 0 | 0.0731817 |
| #4 | Hallmark.GseaPreranked.1585308626238 | HALLMARK_P53_PATHWAY | 2.4347882 | 0 | 0.0025373 |
| #5 | CanonicalPathways.GseaPreranked.1585307356087 | KEGG_GLYCOSAMINOGLYCAN_DEGRADATION | 2.4102724 | 0 | 0.0635057 |
| #6 | OncogenicSignatures.GseaPreranked.1585307079701 | RB_P130_DN.V1_DN | 2.31935 | 0.0019881 | 0.0192678 |
| #7 | CanonicalPathways.GseaPreranked.1585307356087 | PID_P53DOWNSTREAMPATHWAY | 2.2453642 | 0 | 0.1249576 |
| #8 | OncogenicSignatures.GseaPreranked.1585307079701 | TGFB_UP.V1_UP | 2.224309 | 0.0040733 | 0.0230541 |
| #9 | CanonicalPathways.GseaPreranked.1585307356087 | PID_INTEGRIN_CS_PATHWAY | 2.2107453 | 0 | 0.1158633 |
| #10 | CanonicalPathways.GseaPreranked.1585307356087 | REACTOME_EICOSANOID_LIGAND_BINDING_RECEPTORS | 2.2100172 | 0.0018868 | 0.0933814 |
| #11 | CanonicalPathways.GseaPreranked.1585307356087 | REACTOME_INTERFERON_GAMMA_SIGNALING | 2.207767 | 0.0019685 | 0.0790613 |
| #12 | CanonicalPathways.GseaPreranked.1585307356087 | PID_CDC42_PATHWAY | 2.191459 | 0 | 0.0756438 |
| #13 | CanonicalPathways.GseaPreranked.1585307356087 | PID_A6B1_A6B4_INTEGRIN_PATHWAY | 2.1810114 | 0 | 0.0719648 |
| #14 | CanonicalPathways.GseaPreranked.1585307356087 | REACTOME_GENERIC_TRANSCRIPTION_PATHWAY | 2.1670666 | 0 | 0.0712161 |
| #15 | OncogenicSignatures.GseaPreranked.1585307079701 | PRC2_EDD_UP.V1_DN | 2.127989 | 0.0020619 | 0.0346475 |
| #16 | CanonicalPathways.GseaPreranked.1585307356087 | REACTOME_ADHERENS_JUNCTIONS_INTERACTIONS | 2.1265082 | 0.0079681 | 0.082924 |
| #17 | Hallmark.GseaPreranked.1585308626238 | HALLMARK_HEME_METABOLISM | 2.11698 | 0.002079 | 0.0231103 |
| #18 | Hallmark.GseaPreranked.1585308626238 | HALLMARK_REACTIVE_OXYGEN_SPECIES_PATHWAY | 2.1090298 | 0.0020243 | 0.016187 |
| #19 | CanonicalPathways.GseaPreranked.1585307356087 | BIOCARTA_DC_PATHWAY | 2.102159 | 0 | 0.0872241 |
| #20 | CanonicalPathways.GseaPreranked.1585307356087 | PID_INTEGRIN1_PATHWAY | 2.088648 | 0.003937 | 0.0862467 |
| #21 | CanonicalPathways.GseaPreranked.1585307356087 | BIOCARTA_CYTOKINE_PATHWAY | 2.0693016 | 0 | 0.0900062 |
| #22 | CanonicalPathways.GseaPreranked.1585307356087 | ST_P38_MAPK_PATHWAY | 2.0198488 | 0.004 | 0.1135944 |
| #23 | CanonicalPathways.GseaPreranked.1585307356087 | KEGG_OLFACTORY_TRANSDUCTION | 1.9949967 | 0.0040241 | 0.1220827 |
| #24 | CanonicalPathways.GseaPreranked.1585307356087 | REACTOME_G_ALPHA1213_SIGNALLING_EVENTS | 1.9836398 | 0.0019157 | 0.1218731 |
| #25 | CanonicalPathways.GseaPreranked.1585307356087 | PID_MET_PATHWAY | 1.9831402 | 0.0039216 | 0.1151159 |

**Table 2. Top 25 upregulated and downregulated signatures in relapsed versus resected brain metastasis transcriptome.**
| ID path way | Dataset | Pathway | NES | Nominal p-value | FDR q-value |
| --- | --- | --- | --- | --- | --- |
| #1 | Hallmark.GseaP reranked.1585308626238 | HALLMARK_MYC_TARGETS_V1 | -7.075377 | 0 | 0 |
| #2 | Hallmark.GseaP reranked.1585308626238 | HALLMARK_E2F_TARGETS | -6.761198 | 0 | 0 |
| #3 | CanonicalPathways.GseaPreranked.1585307356087 | REACTOME_CELL_CYCLE | -6.609566 | 0 | 0 |
| #4 | CanonicalPathways.GseaPreranked.1585307356087 | REACTOME_CELL_CYCLE_MITOTIC | -6.090444 | 0 | 0 |
| #5 | CanonicalPathways.GseaPreranked.1585307356087 | REACTOME_METABOLISM_OF_RNA | -5.91368 | 0 | 0 |
| #6 | CanonicalPathways.GseaPreranked.1585307356087 | REACTOME_METABOLISM_OF_MRNA | -5.440457 | 0 | 0 |
| #7 |  | CO-OPTION DOWN GENES LOG2FC_1 | -5.316188 | 0 | 0 |
| #8 | CanonicalPathways.GseaPreranked.1585307356087 | REACTOME_METABOLISM_OF_PROTEINS | -5.30438 | 0 | 0 |
| #9 | CanonicalPathways.GseaPreranked.1585307356087 | REACTOME_DNA_REPLICATION | -5.185881 | 0 | 0 |
| #10 | CanonicalPathways.GseaPreranked.1585307356087 | REACTOME_MITOTIC_M_M_G1_PHASES | -5.062673 | 0 | 0 |
| #11 | Hallmark.GseaP reranked.1585308626238 | HALLMARK_OXIDATIVE_PHOSPHORYLATION | -5.022407 | 0 | 0 |
| #12 | Hallmark.GseaP reranked.1585308626238 | HALLMARK_G2M_CHECKPOINT | -4.986867 | 0 | 0 |
| #13 | CanonicalPathways.GseaPreranked.1585307356087 | REACTOME_INFLUENZA_LIFE_CYCLE | -4.916678 | 0 | 0 |
| #14 | CanonicalPathways.GseaPreranked.1585307356087 | REACTOME_TRANSLATION | -4.904846 | 0 | 0 |
| #15 | CanonicalPathways.GseaPranked.1585307356087 | KEGG_RIBOSOME | -4.633262 | 0 | 0 |
| #16 | CanonicalPathways.GseaPranked.1585307356087 | REACTOME_3_UTR_MEDIATED_TRANSLATIONAL_REGULATION | -4.455745 | 0 | 0 |
| #17 | CanonicalPathways.GseaPranked.1585307356087 | REACTOME_MRNA_PROCESSING | -4.444215 | 0 | 0 |
| #18 | CanonicalPathways.GseaPranked.1585307356087 | REACTOME_NONSENSE_MEDIATED_DECAY_ENHANCED_BY_THE_EXON_JUNCTION_COMPLEX | -4.444211 | 0 | 0 |
| #19 | CanonicalPathways.GseaPranked.1585307356087 | REACTOME_INFLUENZA_VIRAL_RNA_TRANSCRIPTION_AND_REPLICATION | -4.390086 | 0 | 0 |
| #20 | CanonicalPathways.GseaPranked.1585307356087 | REACTOME_IMMUNE_SYSTEM | -4.385225 | 0 | 0 |
| #21 | CanonicalPathways.GseaPranked.1585307356087 | REACTOME_PROCESSING_OF_CAPPED_INTRON_CONTAINING_PRE_MRNA | -4.311844 | 0 | 0 |
| #22 | CanonicalPathways.GseaPranked.1585307356087 | KEGG_SPLICEOSOME | -4.30086 | 0 | 0 |
| #23 | CanonicalPathways.GseaPranked.1585307356087 | REACTOME_SRP_DEPENDENT_COTRANSLATIONAL_PROTEIN_TARGETING_TO_MEMBRANE | -4.285922 | 0 | 0 |
| #24 | CanonicalPathways.GseaPranked.1585307356087 | REACTOME_ADAPTIVE_IMMUNE_SYSTEM | -4.253354 | 0 | 0 |
| #25 | CanonicalPathways.GseaPranked.1585307356087 | REACTOME_MITOTIC_G1_G1_S_PHASES | -4.227897 | 0 | 0 |

Based on these results, and given the ability of DEBIO-0932 to effectively target the early stages of metastasis *ex vivo* (Fig. 1, E-G) and *in vivo* (Fig. S3, N-S), we used the HSP90 inhibitor in an adjuvant setting after neurosurgery. DEBIO-0932 administration at 160mg/kg starting 3 days after surgery and maintained for 5-6 weeks efficiently delayed local relapse (Fig. 3, H-J). Interestingly, animals relapsed approximately 2 weeks after treatment was discontinued, with one animal remaining with stable disease over 18 weeks (Fig. 3 J). Given the significant delay in relapse observed during the treatment period and the correlation of metastasis re-growth after DEBIO-0932 was discontinued, maintenance of DEBIO-0932 treatment is expected to improve the survival benefits for longer period of time (Fig S3 J).

These results validate METPlatform as an effective *ex vivo* drug-screening tool for the identification of brain metastasis vulnerabilities, such as HSP90, that could be translated to *in vivo* metastasis assays. In this sense, our findings suggest that inhibition of HSP90 could become a novel strategy to target established brain metastases and/or to prevent their relapse after neurosurgery.

### Patient-derived organotypic cultures (PDOC) are sensitive to HSP90 inhibition independently of the primary tumor origin and established oncogenic drivers

Among the many advantages of METPlatform, the possibility of adapting it to patient-derived organotypic cultures (PDOCs) using fresh surgically-resected human tissue is invaluable for translational purposes.

Brain metastasis PDOCs (BrM-PDOCs) were established from neurosurgical resections performed on ten patients diagnosed with four different types of primary tumor and oncogenic profiles (Fig. 4, A, D and E). Remarkably, DEBIO-0932 treatment, at doses compatible with levels detected in mouse brains with metastases (Fig. 1 J), blunted tumor proliferation in all BrM-PDOCs (Fig. 4, B and C) independently of their primary origin (Fig. 4, D).

**Figure 4.**
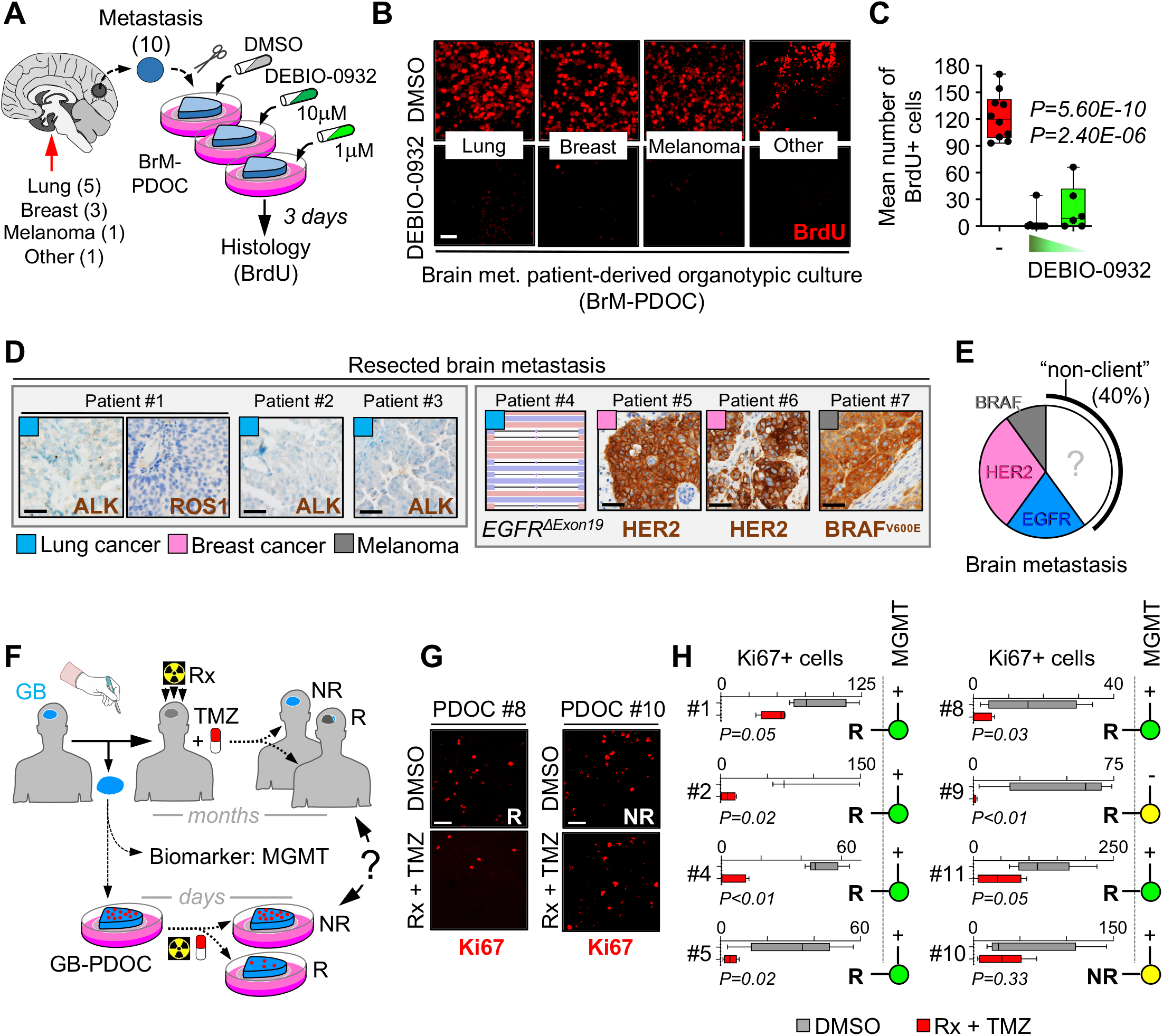
Patient-derived organotypic cultures (PDOC) are sensitive to HSP90 inhibition independently of the primary tumor origin and established oncogenic drivers. **(A)** Schema of the experimental design. Fresh surgically-resected human brain metastases (n=10) from various primary origins were used to perform patient-derived organotypic cultures (BrM-PDOCs) and treated with DEBIO-0932 at 10 μM and 1 μM for 3 days. **(B)** Representative BrM-PDOCs from lung (n=5), breast (n=3), melanoma (n=1) and another primary source (n=1) stained against BrdU. Scale bar, 50 μm. **(C)** Quantification of number of BrdU+ cancer cells found in DMSO and DEBIO-0932 treated BrM-PDOCs. Values are shown in box-and-whisker plots where every dot represents a different patient (mean value obtained from all PDOCs from the same condition and patient) and the line in the box corresponds to the median. Whiskers go from the minimum to the maximum value (n=10 patients with DMSO treated PDOCs, n=9 DEBIO-0932 10 μM and n=6 DeBIO-0932 1 μM, each patient is an independent experiment). *P* value was calculated using two-tailed t-test. **(D)** Representative images of human brain metastases from which BrM-PDOC were generated (AC) showing the status of the HSP90-dependent oncogenic drivers ALK, ROS1, HER2 and BRAF. Scale bar, 50 μm. Targeting sequencing of the *EGFR* locus of a lung cancer brain metastasis patient showing a deletion in exon 19 is also shown. **(E)** Pie chart showing the distribution of the ten BrM-PDOCs with oncogenic drivers sensitive to HSP90 inhibition (Non-HSP90 client: n=4; *EGFR* mutant lung cancer: n=2; HER2+ breast cancer: n=3; *BRAF* mutant melanoma: n=1). **(F)** Schema of experimental design using GB-PDOCs. **(G)** Representative GB-PDOCs responding or not to the standard of care that was provided *ex vivo* (Radiation (Rx): 2x 10 Gy + temozolomide (TMZ) 250 μM). GB organotypic cultures were stained with proliferation markers. NR: nonresponder, R: responder. Scale bar, 50 μm. **(H)** Quantification of number of proliferative cancer cells found in DMSO and Rx + TMZ treated GB-PDOCs. Each patient is represented in an individual graph. Values are shown in box- and-whisker plots and the line in the box corresponds to the median. Whiskers go from the minimum to the maximum value (n=3-6 PDOCs per experimental condition. Eight different patients included, each patient is an independent experiment). *P* value was calculated using two-tailed t-test. MGMT promoter methylation status is shown for each patient (+: methylated; -: unmethylated). A colored circle indicates whether treatment of GB-PDOC correlates with MGMT methylation (green) or not (yellow).

Clinical response to HSP90 inhibitors has been attributed to “addiction” of tumors to particular oncogenes, such as *HER2, ALK, ROS1, EGFR* and *BRAF*, which are sensitive HSP90 client proteins (Neckers and Workman, 2012). Among the five brain metastases from NSCLC, two of them harbored a mutation/deletion in *EGFR* as detected by targeted sequencing (Fig. 4 D and data not shown); however, no molecular alterations in *EGFR, ALK* and *ROS1* were found in the other three patients using standard methodologies approved in clinical practice for these biomarkers (Fig. 4 D). Three brain metastases were derived from HER2+ breast adenocarcinomas, one from a melanoma with the activating mutation BRAF V600E, and one from a gastro-esophageal cancer without known oncogenic drivers sensitive to HSP90 inhibition (Fig. 4 D and data not shown).

Altogether, our results show that METPlatform could be used to validate experimental therapeutic strategies in human samples, where DEBIO-0932 impairs the viability of BrM-PDOCs independently of their primary tumor origin and established HSP90-dependent oncogenes routinely scored in the clinical practice.

However, a major benefit of METPlatform would derive from its use as a strategy for personalized medicine, for instance by providing a fast readout on the efficacy of postsurgical adjuvant treatments. To evaluate PDOCs as *ex vivo* “avatars” of cancer patients we performed a proof-of-concept experiment with glioblastoma (GB) diagnosed *de novo*. In contrast to the lack of standard of care after neurosurgery in patients with brain metastasis, those with GB invariably receive radiotherapy plus temozolomide (Fig. 4 F) (Stupp et al., 2005, 2009). We treated eight GB-PDOCs with the standard of care (radiotherapy and temozolomide) during seven days. The impact of therapy on cancer cell proliferation was compared with MGMT methylation status, an established predictive biomarker of response to this therapy (Hegi et al., 2005; Stupp et al., 2005, 2009), using methylation-specific PCR (Fig. 4, G and H) on the surgical specimen. 6 out of 8 (75%) GB-PDOCs showed the expected response as predicted by the validated biomarker (Fig 4, G and H). However, an unmethylated GB (Patient#9) generated a responsive GB-PDOC and a methylated GB (Patient#10) did not respond when the PDOC was treated (Fig. 4, G and H). With this proof-of-concept, we propose that METPlatform could be used for improving available biomarkers in difficult-to-treat cancer towards optimal patient stratification (Butler et al., 2020).

Consequently, METPlatform shows potential predictive value for therapeutic response and thus should be further evaluated in clinical trials aimed to improve personalized cancer care.

### *In situ* proteomics uncovers HSP90-dependent brain metastasis mediators

Our data support HSP90 as an actionable mediator of brain metastasis. Therefore, we wanted to investigate its biological relevance in this specific progression of cancer since it remains particularly understudied at the molecular level. To identify acute biological responses following HSP90 inhibition, we treated organotypic cultures with established H2030-BrM brain metastases with DEBIO-0932 at 1μM for 6 hours, followed by laser capture microdissection of paraffin-embedded metastatic lesions and peptides identification by mass spectrometry (Fig. 5, A and C). Short time treatment with DEBIO-0932 showed modest but statistically significant reduction of brain metastases as measured by BLI (Fig. 5 B), allowing us to assess early changes after HSP90 inhibition.

**Figure 5.**
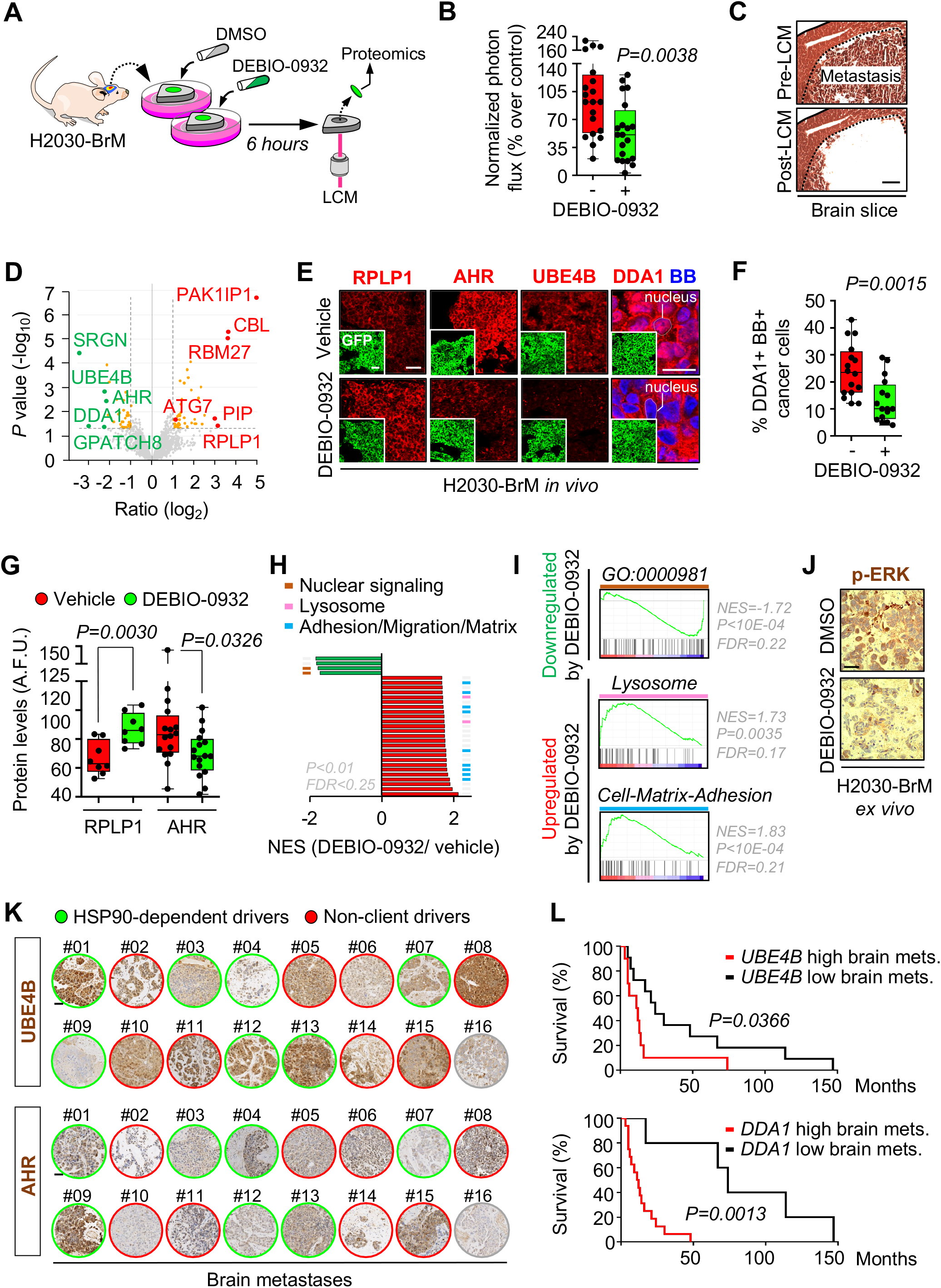
*In situ* proteomics uncovers HSP90-dependent brain metastasis mediators. **(A)** Schema of experimental design. **(B)** Quantification of bioluminescence emitted by H2030-BrM established metastases in brain organotypic cultures. BLI values were obtained 6h after the addition of DEBIO-0932 or DMSO normalized by the values of each culture before any treatment. Data is shown as relative to DMSO BLI values in box-and-whisker plots where every dot represents an organotypic culture and the line in the box corresponds to the median. Whiskers go from the minimum to the maximum value (n=20 organotypic cultures per experimental condition, 3 independent experiments). *P* value was calculated using two-tailed t-test. **(C)** Representative image of a fully established brain metastasis from H2030-BrM before and after laser capture microdissection (LCM). The dotted line delimits the metastasis. Scale bar, 100 μm. **(D)** Volcano plot with deregulated proteins (red: upregulated; green: downregulated) found in brain metastases treated with DEBIO-0932 compared to DMSO (n=3 biological replicates (mice) per condition, n≥12 brain metastases per mouse were pooled together). Proteins with a *P* value <0.05 and a log_2_ ratio >1 or <-1 were defined as deregulated. Gray dotted lines indicate *P* value and log2 ratio cut offs. The names of the top deregulated proteins are shown. **(E)** Representative images showing RPLP1, AHR, UBE4B and DDA1 levels in brain metastases (generated by intracardiac inoculation of H2030-BrM) found at endpoint of vehicle and DEBIO-0932 treated animals. This result was reproduced in 2 independent staining with different brains. BB: bisbenzamide. Scale bars, 50 μm; high magnification, 12 μm. **(F)** Quantification of percentage of nuclear DDA1+ BB+ cells shown in (E). Values are shown in box-and-whisker plots where each dot is a metastatic lesion and the line in the box corresponds to the median. Whiskers go from the minimum to the maximum value (n=16 metastatic lesions from 4 brains per condition, 2 independent staining with different brains were performed). *P* value was calculated using two-tailed t-test. **(G)** Quantification of RPLP1 and AHR levels shown in (E) in arbitrary fluorescent units (A.F.U.). Values are shown in box-and-whisker plots where each dot is a metastatic lesion and the line in the box corresponds to the median. Whiskers go from the minimum to the maximum value (n=8-16 metastatic lesions from 2-4 brains per condition, 2 independent staining with different brains were performed). P value was calculated using two-tailed t-test. **(H)** GSEA of top 25 upregulated (red) and downregulated (green; only four fulfill the filter) pathways upon DEBIO-0932 treatment. Those biological processes represented with more than one signature are labelled with colored lines. **(I)** Examples of signatures included in the main biological processes represented in the proteomic analysis. GO:0000981 corresponds to the Gene Ontology signature “DNA binding, Transcription factor activity, RNA polymerase II specific”. **(J)** Representative images showing p-ERK levels in organotypic cultures from (B). This result was reproduced in 3 independent staining with organotypic cultures from different mice. Scale bar, 20 μm. **(K)** Immunohistochemistry against UBE4B and AHR in 16 human brain metastases with (green) and without (red) HSP90-dependent oncogenic drivers. No molecular information available for the sample depicted in gray. Scale bar, 50 μm. **(L)** Kaplan-Meier curves showing significant correlation between worse survival post-brain metastasis (SPBM) and high expression levels of *UBE4B* (upper panel) or *DDA1* (lower panel) in a cohort of 21 breast cancer brain metastasis patients.

We identified 83 significantly deregulated proteins upon treatment with DEBIO-0932, from which 44 were upregulated and 39 were downregulated (Fig. 5 D). We validated this analysis with immunofluorescence applied to brains from mice treated with DEBIO-0932 *in vivo* to score top deregulated proteins. Increased levels of 60S acidic ribosomal protein P1 (RPLP1) as well as the reduced levels of aryl hydrocarbon receptor (AHR), the ubiquitin conjugation factor E4 B (UBE4B) and the DET1- and DDB1-associated protein 1 (DDA1) (Fig. 5, E-G), whose reduction was mainly observed in the nuclear compartment (Fig. 5, E and F), were all confirmed.

Downregulated proteins upon DEBIO-0932 treatment are potential HSP90-dependent mediators of brain metastasis (Fig. 5 D). Surprisingly, among the top five downregulated proteins (Fig. 5 D), only AHR has been functionally linked to HSP90 (Chen et al., 2013), which could suggest the presence of HSP90-related biological programs independent of the well-established cancer-related HSP90 clients (Whitesell and Lindquist, 2005). In addition, four top downregulated proteins (AHR, UBE4B, DDA1 and GPATCH8 (G patch domain-containing protein 8)) have been shown to be able to translocate into the nucleus (Cheng et al., 2017; Du et al., 2016; Murray et al., 2014) and 50% of top downregulated signatures belong to nuclear signaling pathways (Fig. 5, H and I, Table 3). The nuclear role of HSP90, including modulation of transcription, chromatin remodeling and DNA damage, has been described in several studies (Antonova et al., 2019; Trepel et al., 2010). Added to our previous findings (Fig. 2, I-K), our results suggest a prominent role of HSP90 or HSP90-dependent downstream program in the nucleus of secondary brain tumors, which remain pending for functional characterization. Nonetheless, we do not rule out the impact of DEBIO-0932 on cytoplasmic HSP90 clients, including cancer-related kinases, in brain metastasis. In fact, a reduction in phosphorylated ERK1/2 (Thr202/Tyr204) was detected in organotypic cultures with established brain metastases treated with DEBIO-0932 (Fig. 5 J) in line with previously reported studies (Bao et al., 2009). Thus, we hypothesize that the short period of treatment (Fig. 5 A) limits the detection of significant changes in well-established HSP90 clients (Echeverría et al., 2011) while benefiting others, preferentially nuclear, that could contribute to explain the implication of HSP90 in metastasis.

**Table 3.**
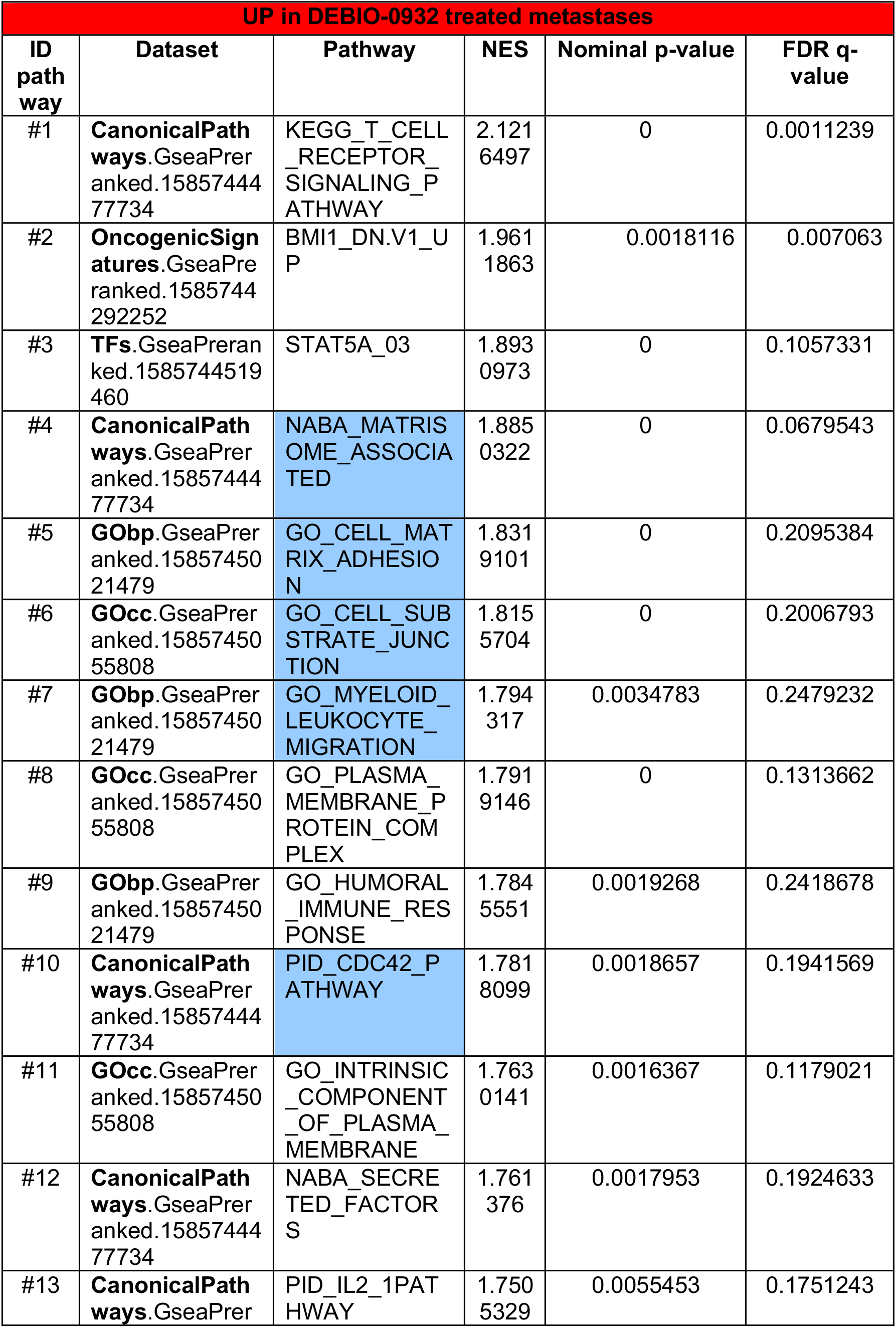
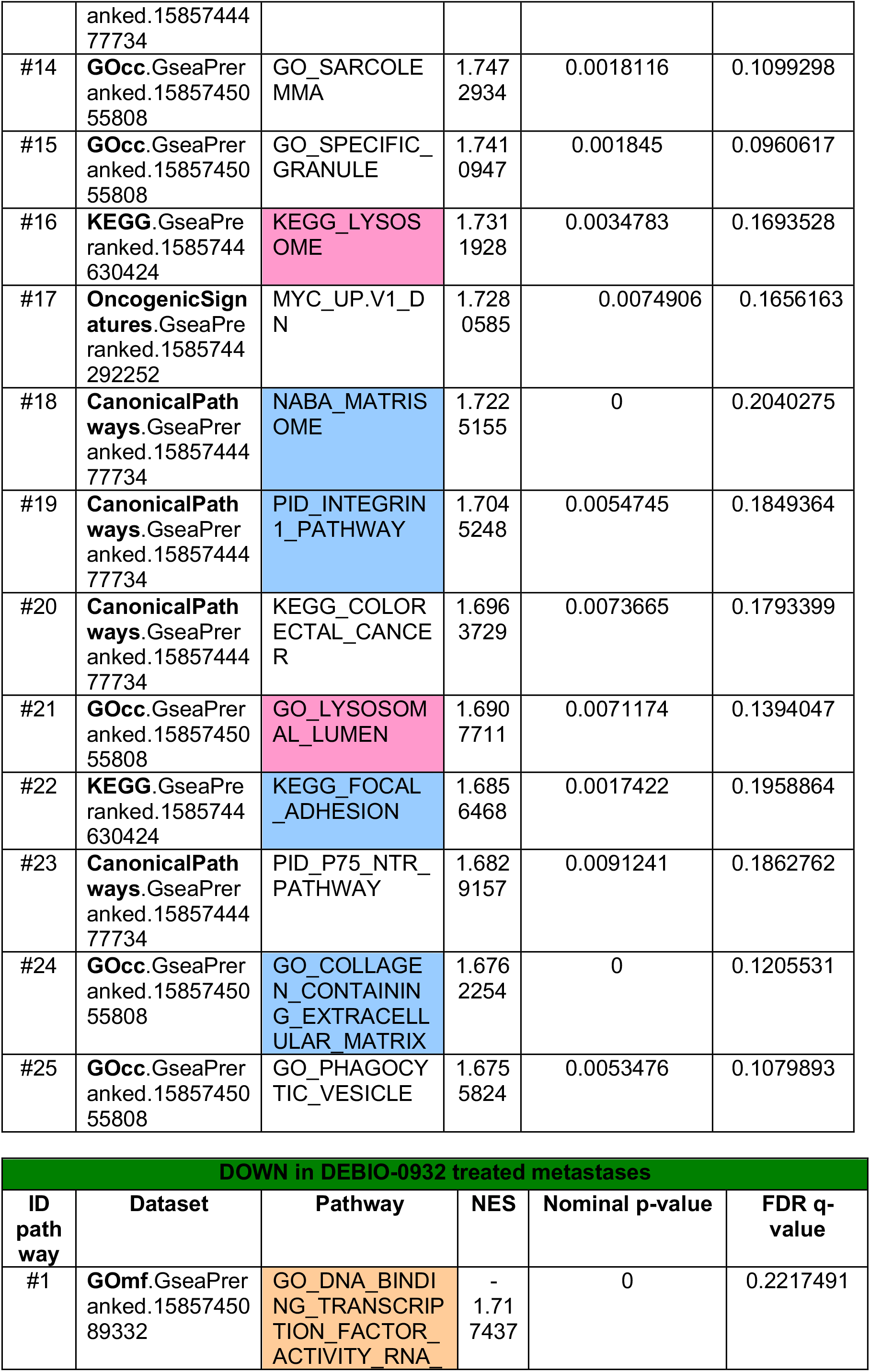
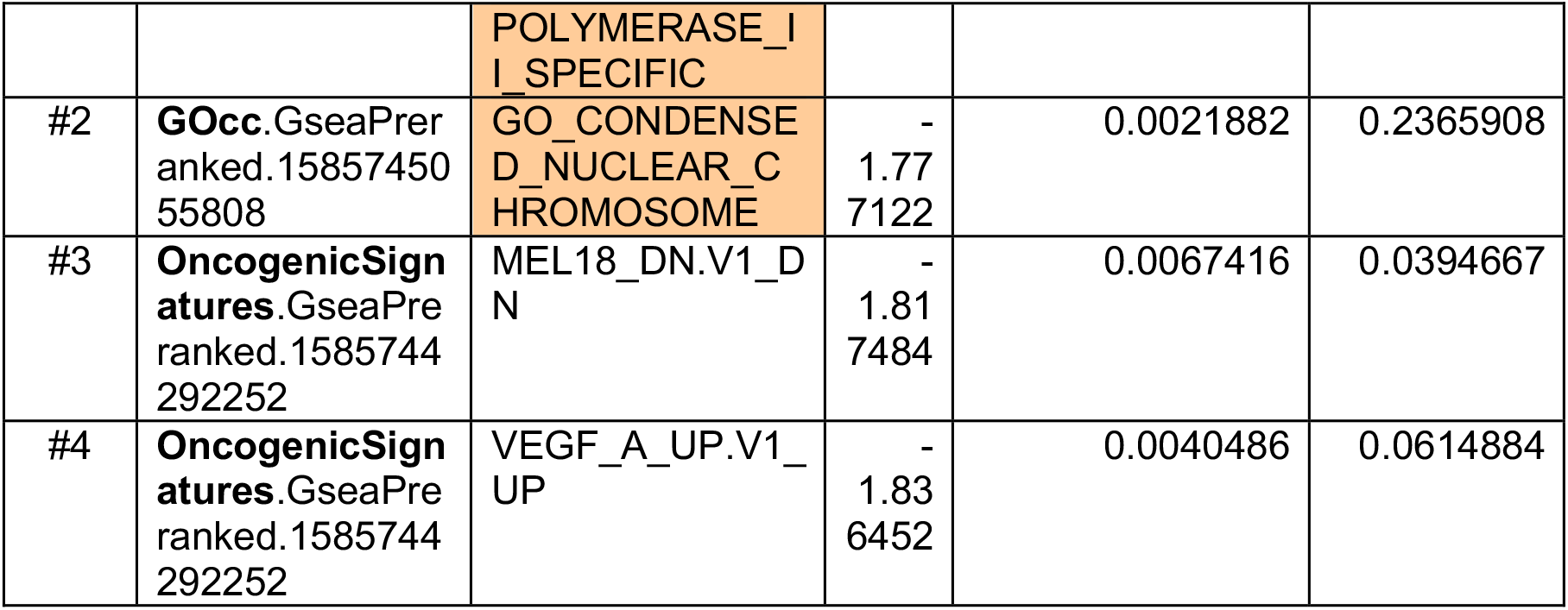
Top 25 upregulated and downregulated signatures in DEBIO-0932 treated lung adenocarcinoma (H2030-BrM) established metastasis. GSEA analysis has been applied to the differentially expressed proteome upon acute DEBIO-0932 treatment of established metastases. Blue: signatures belonging to migration, adhesion and/or matrix. Pink: signatures belonging to lysosome. Yellow: signatures belonging to vascular co-option (*). Dark blue: signatures belonging to cytokine signaling. Light brown: signatures belonging to nuclear signaling.

To further investigate these candidates as novel brain metastasis mediators, we evaluated sixteen human brain metastases by immunohistochemistry and detected UBE4B and AHR expression in all of them independently of the oncogenic driver (Fig. 5 K, Table 4), thus reinforcing our hypothesis about the presence of non-canonical HSP90 downstream programs in brain metastasis. More importantly, *UBE4B* and *DDA1* expression levels in brain metastases were independent prognostic factors of worse survival from the diagnosis of CNS secondary tumors (Fig. 5 L) in a previously published cohort of breast cancer patients (Varešlija et al., 2019). Although validation of these findings in larger and independent cohorts are needed, our results uncover potential brain metastasis biomarkers.

**Table 4.** Molecular profile of human brain metastases. Sixteen brain metastases from 4 different types of cancer and different oncogenomic profiles were analyzed by immunohistochemistry to score HSP90-dependent molecules: UBE4B and AHR.

| Sample ID | Primary tumor | Molecular profile brain metastasis | Immunohistochemistry score |  |
| --- | --- | --- | --- | --- |
|  |  |  | UBE4B | AHR |
| #1 | Lung | EGFR L861Q mutation | 3 | 2 |
| #2 | Lung | ALK-/ ROS1-/ EGFR WT | 2 | 1 |
| #3 | Melanoma | BRAF V600E mutation | 1 | 1 |
| #4 | Breast | HER2+/ ER-/ PR- | 1 | 2 |
| #5 | Melanoma | BRAF WT | 2 | 2 |
| #6 | Lung | ALK-/ ROS1- | 1 | 2 |
| #7 | Breast | HER2+/ ER+/ PR+ | 2 | 1 |
| #8 | Breast | HER2-/ ER-/ PR- | 3 | 2 |
| #9 | Lung | KRAS G12C mutation | 1 | 3 |
| #10 | Lung | ALK-/ ROS1-/ EGFR WT | 2 | 1 |
| #11 | Breast | HER2-/ ER-/ PR- | 3 | 2 |
| #12 | Breast | HER2+/ ER+ | 2 | 1 |
| #13 | Lung | KRAS G12C mutation | 2 | 2 |
| #14 | Colon | KRAS WT | 2 | 2 |
| #15 | Lung | ALK-/ ROS1-/ EGFR WT | 2 | 2 |
| #16 | Potential lung | N/A | 1 | 1 |

Overall, *in situ* proteomic characterization of the *ex vivo* acute response to DEBIO-0932 treatment uncover potential HSP90-dependent mediators, which proves the potential of combining METPlatform with unbiased omic approaches to interrogate brain metastasis.

### METPlatform facilitates unbiased identification of synergistic drug combinations against brain metastasis

Despite the encouraging pharmacological results obtained with DEBIO-0932 *in vivo* (Fig. 3, B-G, Fig. S3, E and F), control of metastatic disease could still be improved. We hypothesized that cancer cells might partially buffer DEBIO-0932-induced cellular toxicity through different resistance mechanisms. By uncovering a potential resistance mechanism to HSP90 inhibitors, our intention is to provide a more solid therapeutic strategy that could eventually be translated to a clinical scenario, where failed efforts to establish HSP90 inhibitors in oncology would benefit from strategies decreasing the therapeutic dose in patients to avoid toxicity (Neckers and Workman, 2012).

Our proteomic analysis on DEBIO-0932 treatment identified the upregulation of multiple signatures representing adhesion, migration and interaction with the matrix as well as increased lysosome activity (Fig. 5, H and I, Table 3), all of which are known mechanisms involved in therapeutic resistance (Orgaz et al., 2020; Sui et al., 2013). Given that lysosome activity is tightly linked to autophagy (Sui et al., 2013) and previous studies reported the induction of autophagy by HSP90 inhibitors in cancer (He et al., 2016; Liu et al., 2012; Mori et al., 2015; Samarasinghe et al., 2014; Zhao et al., 2019), we decided to explore this process as a potential actionable resistance mechanism to HSP90 inhibition in brain metastasis. In addition to the upregulation of the autophagy-related protein ATG7 (Levy et al., 2017) (Fig. 5 D), we noticed that the early response of cancer cells to HSP90 inhibition induced the accumulation of the adaptor protein p62 or sequestosome-1 (Fig. 6, A and B). As an additional evidence of the molecular crosstalk between HSP90 and autophagy in brain metastasis, we used a probe that labels the flux of lysosomal degradation based on GFP-tagged LC3 (Kaizuka et al., 2016). Given the unavailability of H2030-BrM or MDA231-BrM cell lines lacking the GFP reporter (Bos et al., 2009; Chen et al., 2016; Nguyen et al., 2009; Valiente, 2020; Valiente et al., 2014) and that DEBIO-0932 efficacy on brain metastasis is independent of the primary source (Fig. 1, E-I, Fig. S1 G, Fig. 4 B), we used the fluorescence-free melanoma brain metastatic cell line B16/F10-BrM (Priego et al., 2018) for this purpose (Fig. 6 C). Treatment of B16/F10-BrM organotypic brain cultures with DEBIO-0932 decreased the amount of GFP-LC3+ vesicles, which indicates enhanced autophagic flux (Fig. 6, D and E). Note that the same probe also encodes an autophagy-independent RFP reporter, which does not change in the presence of DEBIO-0932 (Fig. 6 D).

**Figure 6.**
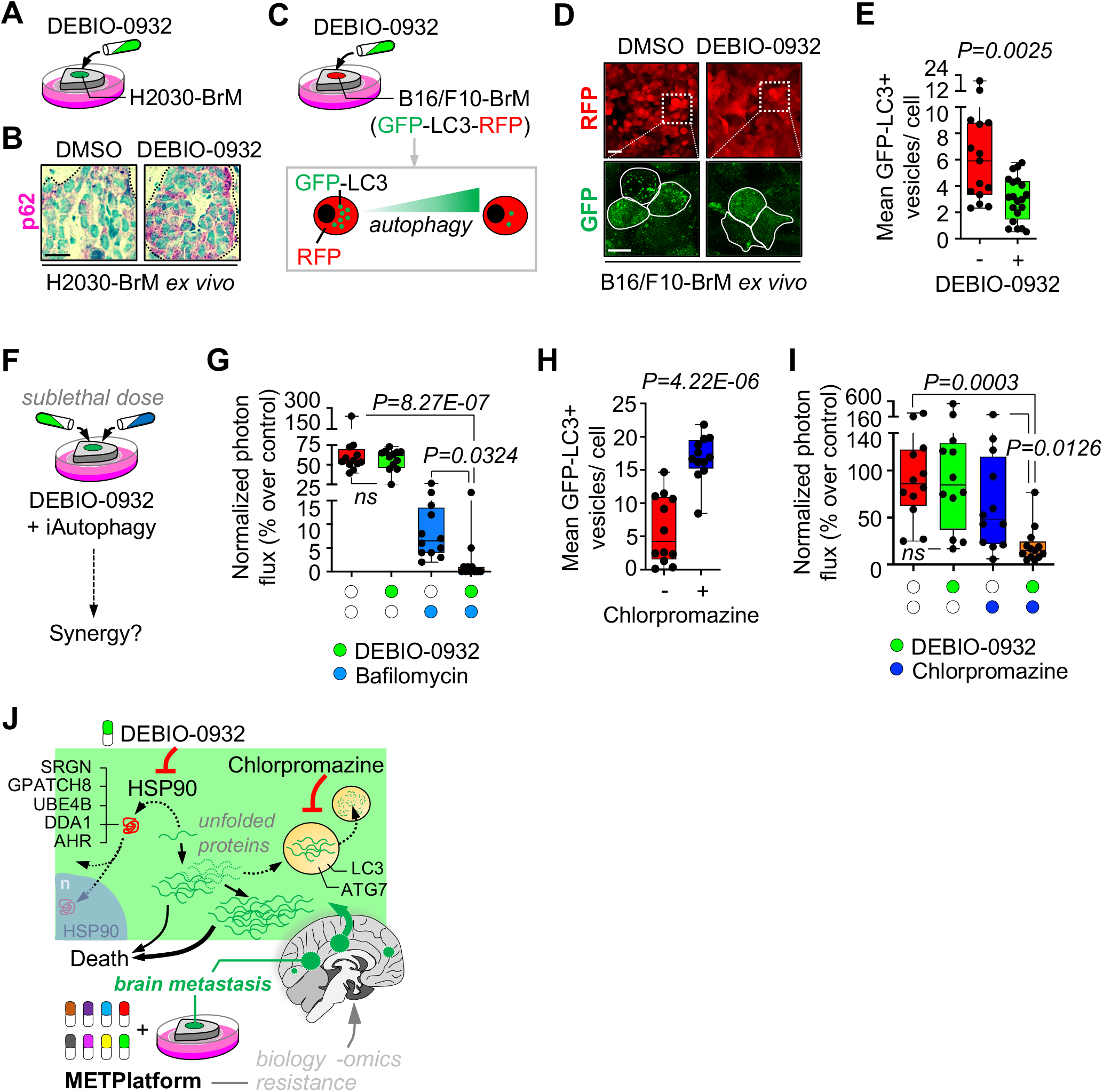
METPlatform facilitates unbiased identification of synergistic drug combinations against brain metastasis. **(A)** Schema of experimental design. Organotypic cultures with established brain metastases from H2030-BrM cells were treated with DEBIO-0932 and evaluated for p62 levels. **(B)** Representative images showing p62 levels. This result was reproduced in 3 independent staining with organotypic cultures from different mice. Dotted lines delimit the metastasis. Scale bar, 10 μm. **(C)** Schema of experimental design. Organotypic cultures with brain metastases from B16/F10-BrM-GFP-LC3-RFP cells were treated with DEBIO-0932 and monitored for autophagic flux by GFP-LC3+ puncta (vesicles). **(D)** Representative organotypic cultures from the experiment in panel (C). RFP is an internal control probe labelling cancer cells independent of autophagy flux and GFP indicate GFP-LC3+ puncta. The dotted line in the upper panel delimits a high magnification area shown in the lower panel respect to the GFP signal derived from GFP-LC3 accumulation. Dotted lines in lower panel surround individual cancer cells. Scale bar, 25 μm; high magnification, 10 μm. **(E)** Quantification of GFP-LC3+ vesicles per cell of the experiment in panel (C). Values are shown in box-and-whisker plots where every dot represents a field of view of an organotypic culture and the line in the box corresponds to the median. Whiskers go from the minimum to the maximum value (DMSO: n=15 fields of view, 2232 cancer cells from 3 organotypic cultures; DEBIO-0932: n=20 fields of view, 3260 cancer cells from 4 organotypic cultures). *P* value was calculated using two-tailed t-test. **(F)** Schema of experimental design. Organotypic cultures with established brain metastases from H2030-BrM cells were treated with DEBIO-0932 and autophagy inhibitors at sublethal doses. **(G)** Quantification of the bioluminescence signal emitted by H2030-BrM cells in each organotypic culture with established brain metastases at day 7 normalized by the initial value at day 0 (before the addition any treatment; both DEBIO-0932 and bafilomycin were added at 100 nM) and normalized to the organotypic cultures treated with DMSO. Values are shown in box-and-whisker plots where every dot represents an organotypic culture and the line in the box corresponds to the median. Whiskers go from the minimum to the maximum value (n=12 organotypic cultures per experimental condition, 2 independent experiments). *P* value was calculated using two-tailed t-test. **(H)** Quantification of GFP-LC3+ vesicles per cell in organotypic cultures with brain metastases from B16/F10-BrM-GFp-LC3-RFP cells treated with chlorpromazine (20 μM) and monitored for autophagic flux by GFP-LC3+ puncta (vesicles). Values are shown in box- and-whisker plots where every dot represents a field of view of an organotypic culture and the line in the box corresponds to the median. Whiskers go from the minimum to the maximum value (DMSO: n=12 fields of view, 1919 cancer cells from 3 organotypic cultures; chlorpromazine: n=12 fields of view, 1759 cancer cells from 3 organotypic cultures). *P* value was calculated using twotailed t-test. **(I)** Quantification of the bioluminescence signal emitted by H2030-BrM cells in each organotypic culture with established brain metastases at day 3 normalized by the initial value at day 0 (before the addition any treatment; DEBIO-0932 was added at 100 nM and chlorpromazine at 20 μM) and normalized to the organotypic cultures treated with DMSO. Values are shown in box-and-whisker plots where every dot represents an organotypic culture and the line in the box corresponds to the median. Whiskers go from the minimum to the maximum value (n=12-13 organotypic cultures per experimental condition, 3 independent experiments). *P* value was calculated using two-tailed t-test. **(J)** Graphical summary. METPlatform is a valuable tool for metastasis research that integrates drug-screening and omic approaches to study pharmacological and biological vulnerabilities. We demonstrate that one vulnerability corresponds to the dependency on HSP90. The BBB-permeable HSP90 inhibitor DEBIO-0932 is an effective therapeutic strategy against established brain metastasis and the analysis of such phenotype with *in situ* proteomics revealed potential novel mediators of brain metastasis downstream HSP90. At the same time, autophagy appears as an actionable mechanism of resistance upon HSP90 inhibition, allowing design of rationale combinations using autophagy inhibitors (i.e. chlorpromazine) and DEBIO-0932 to target brain metastasis more effectively. n: nucleus.

Based on the above findings indicating increased autophagy upon DEBIO-0932 treatment, we combined it with the broadly used autophagy inhibitor bafilomycin A1 (Mauvezin et al., 2015). Combined therapy with both inhibitors in established lung adenocarcinoma H2030-BrM brain metastases *ex vivo* showed synergistic effects compared to sublethal concentration of DEBIO-0932 (Fig. 6, F and G). However, bafilomycin A1 did not progress to clinical development due to its poor toxicity profile *in vivo* (DeVorkin and Lum, 2014; Keeling et al., 1997). Therefore, we looked for alternative compounds able to block autophagy and superior ability to cross the BBB. The FDA-approved antipsychotic drug chlorpromazine fulfills these two requirements (Fig. 6 H) (Nadanaciva et al., 2011). As predicted based on our findings, the combination of sublethal concentration of DEBIO-0932 with the CNS-related drug chlorpromazine was effective against H2030-BrM established brain metastases *ex vivo* (Fig. 6 I).

These results demonstrate the potential of METPlatform not only to uncover previously unrecognized brain metastasis vulnerabilities, but also to exploit them more effectively in combination with unbiased omic approaches. As a result, we report that the BBB-permeable anti-psychotic drug chlorpromazine could be repurposed to target brain metastasis in combination with the HSP90 inhibitor DEBIO-0932 (Fig. 6 J).

## Discussion

The novel drug-screening platform we report here (METPlatform) allows identification of pharmacological vulnerabilities of metastasis *in situ* using an *ex vivo* setting. Organotypic cultures is a well-established technique that could be applied to many different organs (Humpel, 2015; Shamir and Ewald, 2014) and has been reported by us and others to maintain the specific tissue architecture and cellular composition when applied to metastasis (Er et al., 2018; Priego et al., 2018; Valiente et al., 2014; Zhu and Valiente).

Given the vicious cycle (exclusion of patients with CNS disease from clinical trials together with the limited information on the ability of drugs to cross the BBB or blood-tumor barrier (BTB)) that decelerates the development of new therapeutic opportunities for patients with brain metastases, we have applied METPlatform to this clinical need. A hit obtained with METPlatform (i.e. HSP90 inhibitor) was effectively translated *in vivo* using clinically relevant mouse models, including a novel preventive strategy for local relapse, as well as to human brain metastasis.

Although organotypic cultures do have limitations as an experimental drug-screening platform for metastasis (i.e. they do not mimic the whole metastatic cascade or have the ability to score whether a compound crosses a vascular barrier such as the BBB or BTB in the brain), this strategy is highly versatile. For instance, METPlatform is able to resemble different stages of organ colonization and it is compatible with omic approaches, therefore providing a unique tool for basic and translational metastasis research. Future screens applied to brain disorders, including metastasis, might benefit if available artificial intelligence (AI) technology (Saxena et al., 2019) is incorporated to preselect BBB-permeable compounds before applying METPlatform.

Exploiting additional hits scoring *ex vivo* but not *in vitro* are of particular interest since their targets (MEK1/2, CDKs, RAF1, BRAF, VEGFR2, TOP-1, DNA-PK, PI3K, ATR) might be key during organ colonization. Adaptation of cancer cells to a new organ involves molecular changes at various levels including the transcriptome, metabolome or proteome of cancer cells (Basnet et al., 2019; Park et al., 2011; Sevenich et al., 2014). These changes that are only detected *in situ* (i.e. metastatic cells colonizing the brain) could define the presence or absence of a drug target or activate a pathway that drives resistance to a specific therapeutic pressure (Chen et al., 2016). Another category within the hits obtained with METPlatform includes inhibitors that are only effective at advanced stages of the disease. This category might match reported differences within the process of colonization where remodeling of the naïve microenvironment into a protumor niche, where the targets might be present, define early and advanced stages during organ colonization, respectively (Priego et al., 2018; Sevenich et al., 2014). Thus, within this category, anti-metastasis hits could also score by targeting the protumor microenvironment (i.e. pazopanib) (Gril et al., 2013).

Although the association of high HSP90 levels with systemic disease in cancer has been broadly reported (Dimas et al., 2018; Gallegos Ruiz et al., 2008; McCarthy et al., 2008; Pick et al., 2007; Su et al., 2016), whether this association involves any specific step of the metastatic cascade has not been addressed. We demonstrate that the last step of metastasis, organ colonization, is sensitive to HSP90 inhibition, either during the initial stages as well as once metastases are established. Although we focused our efforts to uncover the underlying molecular mechanisms in brain metastasis, the dependency of metastatic colonization on HSP90 does not seem to be organ-specific. However, whether the potential HSP90-regulated proteins (SRGN, DDA1, UBE4B, GPATCH8, AHR) reflect brain-specific functions of the chaperone remains to be determined as well as their specific molecular dependency on HSP90 since, with the exception of AHR (Chen et al., 2013), the rest have not been previously described to be direct substrates. Furthermore, the ability of four out of five identified targets (AHR, DDA1, UBE4B, GPATCH8) to translocate to the nucleus (Cheng et al., 2017; Du et al., 2016; Murray et al., 2014) where they could regulate mechanisms required for brain colonization is currently being investigated.

Second-generation of synthetic HSP90 inhibitors were designed to overcome the pharmacologic limitations of the first ones, showing enhanced potency, reduced toxicity and improved bioavailability, with one compound, DEBIO-0932, even showing BBB penetration (Bao et al., 2009). DEBlO-0932 showed limited clinical efficacy and manageable toxicity in a Phase I clinical trial for patients with advanced solid tumors and lymphoma (Isambert et al., 2015), and is currently under evaluation in a Phase I/II study for patients with advanced NSCLC (NCT01714037), which excludes patients with brain metastases. Recent reports on selected patients enrolled in this trial suggest potential benefit in the metastatic setting (Cedrés et al., 2018), suggesting the possibility that patient stratification could improve the therapeutic benefit. To the best of our knowledge, none of the clinical trials using these inhibitors includes patients with brain metastases. Therefore, our work provides the rationale and proof-of-principle to include patients with CNS disease in current clinical trials with BBB-permeable HSP90 inhibitors.

Our ability to model local relapse after neurosurgery represents not only a clear opportunity to translate our findings into a clinically-compatible preventive strategy, but also allows to investigate the biology of this process for the first time. Our findings on the molecular characterization of relapse suggest that cancer cells left behind after a debulking neurosurgery reinitiate the metastasis following similar mechanism than those needed during metastasis initiation such as vascular co-option (García-Gómez and Valiente, 2020). Exploiting the molecular mechanisms underlying local relapse offers a unique opportunity to study the crosstalk between metastasis initiating cells and a severely damaged microenvironment where the tumor will be regenerated over time.

A more general conclusion of our results is that METPlatform could become a novel tool with direct applications in the clinic. Patient “avatars” have been exploited using patient-derived xenografts (PDXs), patient-derived organoids (PDOs) and patient-derived primary cultures (PDCs) to test candidate drugs prior to the treatment of the patient, therefore helping to select empirical therapies for the study (Gao et al., 2015; Garralda et al., 2014; Lee et al., 2018; Vlachogiannis et al., 2018). Predicting patient response to a specific treatment as early as possible is very relevant in the current context of personalized medicine to optimize the benefits of clinical trials for patients. In this regard, we have confirmed the ability of METPlatform not only to reproduce the correlation between established biomarkers and therapeutic response in difficult-to-treat cancers (i.e. glioblastoma) but also its potential to further dissect existing discordances in the clinical practice.

We are confident that METPlatform could improve the impact of personalized medicine in oncology if applied to patients at high risk of developing resistance or when no biomarker-driven therapies exist, which are aspects of especial relevance in metastasis.

## Acknowledgements

We want to thank all members from the Brain Metastasis Group, F.X. Real, M. Malumbres and M. Barbacid for critical discussion of the manuscript, the CNIO Core Facilities for their excellent assistance, Antonio Cebriá and Javier Klett for their excellent assistance in drug-screening, Sonsoles Rodríguez-Arístegui for her excellent work with preparation of different compounds and Alexandra de Francisco, Yolanda Sierra and María de la Jara Felipe for their excellent work with animal preparation and imaging protocols. We also thank J. Massagué (MSKCC) for some of the BrM cell lines. This work was supported by MINECO (SAF2017-89643-R, SAF2014-57243-R, SAF2015-62547-ERC) (M.V.), Fundación FERO (IX FERO Grant for Research in Oncology) (M.V.), Fundació La Marató de TV3 (141) (M.V.), Melanoma Research Alliance (Bristol-Myers Squibb-Melanoma Research Alliance Young Investigator Award 2017 (498103)) (M.V.), Beug Foundation (Prize for Metastasis Research 2017) (M.V.), Fundación Ramón Areces (CIVP19S8163) (M.V.), Worldwide Cancer Research (19-0177) (M.V.), H2020-FETOPEN (828972) (M.V.), Cancer Research Institute (Clinic and Laboratory Integration Program CRI Award 2018 (54545)) (M.V.), AECC (Coordinated Translational Groups 2017 (GCTRA16015SEOA) (M.V.), LAB AECC 2019 (LABAE19002VALI) (M.V.)), ERC CoG (864759) (M.V.), Sophien-Stiftung zur Förderung der klinischen Krebsorschung (T.W.), Stiftung für angewandte Krebsforschung (T.W.), Forschungskredit of the University of Zurich (FK-18-054) (T.W.), Betty and David Koetser Foundation for Brain Research (T.W.), Comunidad de Madrid (S2017/BMD-3867 RENIM-CM and Y2018/NMT-4949 NanoLiver-CM) and European structural and investment funds (M.D.), La Caixa-Severo Ochoa International PhD Program Fellowship (LCF/BQ/SO16/52270014) (L.Z.), Boehringer Ingelheim Fonds MD fellowship (L.M.). The contribution of the Experimental Therapeutics Programme was supported by core funding from the Spanish National Cancer Research Center (CNIO). CNIO is supported by the ISCIII, the Ministerio de Ciencia e Innovación, and is a Severo Ochoa Center of Excellence (SEV-2015-0510). The CNIC is supported by the ISCIII, the Ministerio de Ciencia e Innovación and the Pro CNIC Foundation, and is a Severo Ochoa Center of Excellence (SEV-2015-0505). M.V. is a Ramón y Cajal Investigator (RYC-2013-13365) and EMBO YIP (4053).

## Author contributions

L.Z. and M.V. designed and performed the experiments, analyzed the data and wrote the manuscript. N.Y. and D.R. performed the experiments and analyzed the data. L.M. analyzed the correlation between gene expression and patient survival. E.H.E. and L.Z. performed the pharmacokinetic experiments and analyzed the data. T.W., J.M., M.W. and L.Z. performed the proteomic experiments and analyzed the data. D.M. quantified the immunofluorescence images. O.G.C. performed the transcriptomic analysis. C.N. and M.D.T. provided reagents and technical expertise with the neurosurgery experiments. L.C. and M.D. performed the MRI experiments and analyzed the data. L.B., P.C., R.S., J.M.S, A.P.N., A.H.L., Y.R. and O.T. provided the human samples and analyzed the clinical data. E.C. scored the clinical samples. J.P. selected and proposed DEBIO-0932 and chlorpromazine as candidates for described experiments. C.B.A. and S.M. performed the *in vitro* screening and provided the chemical inhibitors.

## Methods

### Chemicals and reagents

An in-house chemical library composed of 114 FDA-approved or in clinical trials anti-tumoral drugs (Bejarano et al., 2019) solved in DMSO (Sigma-Aldrich) was used for *ex vivo* and *in vitro* screening. For *in vivo* treatment, DEBIO-0932 (MedChemExpress) was formulated in 30% captisol (Ligand). For *ex vivo* and *in vitro* treatments, DEBIO-0932, bafilomycin A1 (Selleckchem) and chlorpromazine (Sigma-Aldrich) were solved in DMSO.

### Animal studies

All animal experiments were performed in accordance with a protocol approved by the CNIO (IACUC.030-2015), Instituto de Salud Carlos III (CBA35_2015-v2) and Comunidad de Madrid Institutional Animal Care and Use Committee (PROEX250/15 and PROEX135/19). Athymic nu/nu (Harlan) 4-10 weeks of age were used. Brain colonization assays were performed by injecting 100 μL PBS into the left ventricle containing 100,000 cancer cells or 2 μL RPMI1640 intracranially (the right frontal cortex, approximately 1.5 mm lateral and 1 mm caudal flow bregma, and to a depth of 1 mm) containing 100,000 cancer cells by using a gas-tight Hamilton syringe and a stereotactic apparatus. Brain colonization was analyzed *in vivo* and *ex vivo* by BLI. Anesthetized mice (isoflurane) were injected retro-orbitally with D-luciferin (150 mg/kg; Syd Labs) and imaged with an IVIS machine (Perkin Elmer). Bioluminescence analysis was performed using Living Image software, version 4.5. Brain tumor resection was performed by adapting previously described procedures (Morrissy et al., 2016). In brief, after exposing the skull, a craniotomy is performed surrounding the tumor area, which is visualized by GFP. The skull and the dura are lifted with micro-dissecting forceps, the bulk of the tumor is then removed using a microcurette guided by GFP. When hemostasis is obtained, the surgical wound is sutured using interrupted stitching with absorbable sutures. Animals receive meloxicam at 5 mg/kg once per day during 72h and dexamethasone at 13 mg/kg once per day during 48h to contain brain edema. DEBIO-0932 was administered by oral gavage (160 mg/kg) for 3 weeks, daily during the first week and once every 48h during the second week, starting 7 or 14 days after intracardiac inoculation of cancer cells for preventive or interventive therapy, respectively. For preventive therapy of relapse after neurosurgery, DEBIO-0932 was administered by oral gavage (160 mg/kg) for 5-6 weeks, starting 3 days after neurosurgery.

### Organotypic cultures

Organotypic cultures from adult mouse brain and liver were prepared as previously described (Valiente et al., 2014). Brains with established metastases (5-7 weeks after intracardiac inoculation of cancer cells) or without metastases (wild type) and wild type livers were used. In brief, organs were dissected in HBSS supplemented with HEPES (pH 7.4, 2.5 mM), D-glucose (30 mM), CaCl2 (1 mM), MgCl2 (1 mM) and NaHCO3 (4 mM), and embedded in 4% low-melting agarose (Lonza) preheated at 42 °C. The embedded organs were cut into 250 μm slices using a vibratome (Leica). Brain slices were divided at the hemisphere into two pieces. Slices were placed with flat spatulas on top of 0.8 μm pore membranes (Sigma Aldrich) floating on slice culture media (DMEM, supplemented HBSS, FBS 5%, L-glutamine (1 mM) and 100 IU/mL penicillin/streptomycin). Brain slices were imaged to confirm the presence of established metastases using BLI (day 0) and were cultured in the presence of the anti-tumoral library at 10 μM of each compound. Brain slices were imaged 3 days after the addition of the inhibitors (day 3). For 7 days cultures, treatments were replaced at day 3 in fresh media and slices were imaged at day 7. If slices were obtained from wild type brains, 30,000 cancer cells suspended in 2 μL of slice culture media were placed on the surface of the slice and incubated in the presence of the inhibitors for 4 days. Brain slices were imaged 12-16h after addition of cells (day 0) and 3 days after the first BLI (day 3). Growth rate was obtained by comparing the fold increases between day 3/7 and day 0, and normalized to values obtained from slices cultured with DMSO (100%). The BrdU pulse (0.2 mg/mL, Sigma-Aldrich) was given by adding it in the media 2h (H2030-BrM) or 4h (MDA231-BrM) before fixation. Brain slices were fixed in 4% paraformaldehyde (PFA) overnight and then free-floating immunofluorescence was performed. For proteomic analysis, organotypic cultures with established brain metastases from H2030-BrM were treated with DEBIO-0932 at 1 μM for 6h followed by fixation with 4% PFA overnight at 4°C. For analysis of autophagic flux, 200,000 B16/F10-BrM cells with stable expression the autophagy probe GFP-LC3-RFP were added on wild type brain slices and incubated for 24h, followed by DEBIO-0932 treatment at 10 μM for 12h and fixation with 4% PFA overnight at 4°C. For evaluation of hepatotoxicity, wild type liver slices were cultured in the presence of the corresponding inhibitors for 3 days and fixed with 4% PFA overnight at 4°C followed by free-floating immunofluorescence.

### Patient-derived organotypic cultures (PDOCs)

Surgically-resected human brain metastases and newly diagnosed glioblastomas were collected in neurobasal media A supplemented with 1x B27 (17504-044, Gibco), 1x N-2 (17502-048, Gibco), 25 ng/mL bFGF (13256029, Gibco), 100 ng/mL IGF1 (291-G1, R&D Systems), 25 ng/mL EGF (E9644, Sigma Aldrich), 10 ng/mL NRG1-β1/HRG-β1 (396-HB, R&D Systems), 100 IU/mL penicillin/streptomycin and 1 μg/mL amphotericin B. Tissue was embedded in 4% low-melting agarose and 250 μm slices were obtained using a vibratome. Slices from brain metastases were cultured in the presence of DEBIO-0932 at 10 μM and 1 μM for 3 days. Slices from glioblastomas were cultured in the presence of temozolomide (Sigma-Aldrich) at 250 μM for 8 days and received 20 Gy of radiation fractionated in 2 doses of 10 Gy at days 1 and 4. Temozolomide treatment was replaced at day 4 in fresh media. A BrdU pulse (4h) was given at the end of the experiment followed by fixation of slices with 4% PFA overnight at 4°C and free-floating immunofluorescence.

### Cell culture

Human and mouse BrM cell lines have been previously described (Bos et al., 2009; Nguyen et al., 2009; Priego et al., 2018; Valiente et al., 2014). H2030-BrM3 (abbreviated as H2030-BrM) and PC9-BrM3 (abbreviated as PC9-BrM) were cultured in RPMI1640 media supplemented with 10% FBS, 2 mM L-glutamine, 100 IU/mL penicillin/streptomycin and 1 μg/mL amphotericin B. MDA231-BrM2 (abbreviated as MDA231-BrM), ErbB2-BrM2 (abbreviated as ErbB2-BrM), 393N1 and B16/F10-BrM3 (abbreviated as B16/F10-BrM) where cultured in DMEM media supplemented with 10% FBS, 2 mM L-glutamine, 100 IU/mL penicillin/streptomycin and 1 μg/mL amphotericin B. For retrovirus production, HEK293T cells were cultured in DMEM media supplemented with 10% FBS, 2 mM L-glutamine, 100 IU/mL penicillin/streptomycin and 1 μg/mL amphotericin B. For the *in vitro* screening with the anti-tumoral library, H2030-BrM cells were seeded in 96-well microtiter plates at a density of 4,000 cells/well. Cells were incubated for 24h before adding the compounds. Compounds were weighed out and solved in DMSO to a final concentration of 10 mM. From here, a “mother plate” with serial dilutions was prepared at 100x the final concentration in the culture. The final concentration of DMSO in the tissue culture media should not exceed 1%. 2 μL of the compounds were added automatically (Beckman FX 96 tip) to 200 μL media to make it up to the final concentration for each drug. Each concentration was assayed in duplicate. Cells were exposed to the compounds for 72h and then processed for CellTiter-Glo^®^ Luminescent Cell Viability Assay (Promega) readout according to manufacturer’s instructions and read on EndVision (Perkin Elmer). Proliferation rate (%) was calculated by normalizing luminescent values obtained for each compound to values obtained with DMSO (100%).

### Retrovirus production

HEK293T cells at 70% confluence were transfected with 10 μg of pMRX-IP-GFP-LC3-RFP (#84573, Addgene), 5 μg of VSV.G (#14888, Addgene) and 5 μg pCL-Eco (#12371, Addgene) in Opti-MEM with Lipofectamine 2000 (Invitrogen) and incubated at 37°C for 8h. Media was replaced with DMEM supplemented with 10% FBS and 2 mM L-glutamine and virus production was maintained for 48h. Viral supernatant was collected, passed through a 0.45 μm syringe filter and added to B16/F10-BrM cells at 50% confluence in DMEM supplemented with 10% FBS, 2 mM L-glutamine and polybrene (5 μg/mL, Sigma Aldrich). The following day, media was replaced with DMEM supplemented with 10% FBS, 2 mM L-glutamine, 100 IU/mL penicillin/streptomycin and 1 μg/mL amphotericin B. Selection with puromycin (2 μg/mL, Sigma Aldrich) was started 48h after and maintained until complete cell death was observed in the non-infected cancer cells.

### Clinical samples and immunohistochemistry

Sixty brain metastases from lung cancer (40 cases) and breast cancer (20 cases) and thirty matched primary tumors (28 lung tumors and 2 breast tumors) were obtained from University Hospital of Turin to assess HSP90 levels. Twenty-two brain metastases were obtained from Hospital Universitario 12 de Octubre to evaluate UBE4B, AHR, ALK, ROS1, HER2 and BRAF^V600E^ status. All samples followed protocols approved by the Institutional Review Board (IRB) (1-18-01/2016) and the Biobank of the hospital, respectively. Immunohistochemistry against HSP90, UBE4B or AHR was performed at the CNIO Histopathology Core Facility using a standardized automated protocol (Ventana Discovery XT, Roche for HSP90; AS Link, Dako, Agilent for UBE4B and AHR). All reagents, with exception of the primary antibodies, were purchased from Roche and Agilent. For HSP90, antigen retrieval was performed using cell conditioning solution (CC1 mild), followed by endogenous peroxides blocking with inhibitor CM. For UBE4B and AHR, antigen retrieval was performed with the appropriate pH buffer (low or high pH buffer, respectively) and endogenous peroxidase was blocked with peroxide hydrogen at 3%. Slides were incubated with the corresponding primary antibodies as follows: HSP90α/β (clone F-8, 1:3,000; sc-13119, Santa Cruz Biotechnology, 8 min), UBE4B (1:500; ab97697, Abcam) and AHR (1:400; 031714, USBiologicals Life Sciences). After the primary antibody, slides were incubated with the corresponding secondary antibodies and visualization systems (OmniMap for HSP90 and EnVision FLEX+Rabbit Linker for UBE4B and AHR) conjugated with horseradish peroxidase. Immunohistochemical reaction was developed using ChromoMap DAB kit. Nuclei were counterstained with hematoxylin. Finally, the slides were dehydrated, cleared and mounted with a permanent mounting medium for microscopic evaluation. Positive control sections known to be primary antibody positive were included for each staining run. Immunostains were blindly evaluated and scored by a pathologist. Intensity of the staining was evaluated and a representative score (the score covering the largest tumor area) was assigned to each sample. Percentage of cancer cells positive for cytoplasmic HSP90 over total tumor area was quantified. Nuclear HSP90 was scored by quantifying percentage of cancer cells positive for nuclear HSP90 normalized to total tumor. Immunohistochemistry against ALK, ROS1, HER2 and BRAF^V600E^ was performed on 4 μm thick sections of formalin-fixed and paraffin-embedded brain samples at Hospital 12 de Octubre. For immunostaining against ROS1 and BRAF^V600E^, a standardized automated protocol using the Leica Bond Polymer Refining Kit (Leica Bond-III stainer, Leica Biosystems) was performed. Antigen retrieval was performed using 30’ EDTA, pH 9.0, followed by endogenous peroxides blocking with hydrogen peroxide. Slides were incubated with the corresponding primary antibodies (ROS1, 1:200; mAb3287, Cell Signaling; anti-BRAF^V600E^ clone VE1, 1:50; E19294, Spring Bioscience). For immunostaining against ALK, a standardized automated protocol using the OptiView DAB IHC Detection Kit (BenchMark GT automated immunostainer, Ventana, Roche) was performed. Antigen retrieval was performed using CC1 92’, followed by endogenous peroxides blocking. Slides were incubated with the primary antibody (anti-ALK clone D5F3, prediluted; 790-4794, Ventana). After the primary antibodies, slides were incubated with the corresponding secondary antibodies and visualization systems (OptiView DAB for ALK and Leica Bond Polymer Refining Kit for ROS1 and BRAF) conjugated with horseradish peroxidase. Immunohistochemical reaction was developed using DAB. Nuclei were counterstained with hematoxylin. Finally, the slides were dehydrated, cleared and mounted with a permanent mounting medium for microscopic evaluation. For immunostaining against HER2, the Bond Oracle HER2 IHC System (Leica Biosystems) was used.

### Immunofluorescence and immunohistochemistry

Tissue for immunofluorescence was obtained after overnight fixation with 4% PFA at 4°C. Slicing of the brain was done by using a vibratome (Leica) or sliding microtome (Thermo Fisher Scientific). Both types of brain slices (250 μm and 80 μm, respectively) were blocked in 10% NGS, 2% BSA and 0.25% Triton X-100 in PBS for 2h at room temperature (RT). Primary antibodies were incubated overnight at 4°C in the blocking solution and the following day for 30 min at RT. After extensive washing in PBS-Triton 0.25%, the secondary antibody was added in the blocking solution and incubated for 2h. After extensive washing in PBS-Triton 0.25%, nuclei were stained with bis-benzamide (1 mg/mL; Sigma-Aldrich) for 7 min at RT. Brain slices were pre-treated with methanol for 20 min at −20°C before the blocking step for nuclear staining against DDA1. For staining against BrdU, mouse brain slices or PDOCs were treated with HCI 2N 30 min at 37°C, followed by 0.1M borate buffer (pH 8.5) incubation for 10 min at RT. After extensive washing in TBS, slices were blocked in 3% NGS in TBS-Triton 0.25% for 1h at RT and primary antibody was incubated for 72h at 4°C. After extensive washing with TBS-Triton 0.25%, the secondary antibody was incubated in blocking solution for 2h at RT followed by extensive washing with TBS. Primary antibodies: GFP (1:1,000; GFP-1020, Aves Labs), BrdU (1:500; ab6326, Abcam), Ki67 (1:500; ab15580, Abcam), HSP90α/β F-8 (1:500; sc-13119; Santa Cruz Biotechnology), HSP70/ HSC70 W27 (1:500; sc-24; Santa Cruz Biotechnology), AHR (1:300; 31.714.200, US Biological), UBE4B (1:100; ab97697; Abcam), RPLP1 (1:100; HPA003368, Sigma-Aldrich), DDA1 (1:100; 14995-1-AP; ProteinTech), NeuN (1:500; MAB377, Millipore), collagen IV (1:1000; AB756P, Millipore). Secondary antibodies: Alexa-Fluor anti-chicken 488, anti-rabbit 488, anti-rat 555, anti-mouse 555, anti-rabbit 555 (dilution 1:300; Invitrogen). Immunohistochemistry staining against p62 and p-ERK was performed using a standardized automated protocol (Ventana Discovery XT, Roche). Antigen retrieval was performed using cell conditioning solution (CC1 mild), followed by endogenous peroxides blocking with peroxide hydrogen at 3%. Slides were incubated with the corresponding primary antibodies (anti-p62 Ick ligand clone 3/P62 LCK LIGAND, 1:50; 610832, BD Biosciences; phosphop44/42 MAPK (Erk1/2), 1:300; #9101, Cell Signaling). Slides were incubated with the corresponding secondary antibodies and visualization systems (OmniMap) conjugated with horseradish peroxidase. Immunohistochemical reaction was developed using Discovery Purple and ChromoMap DAB kits and nuclei were counterstained with Carazzi’s hematoxylin. Finally, the slides were dehydrated, cleared and mounted with a permanent mounting medium for microscopic evaluation.

### *MGMT* methylation-specific PCR (MSP)

DNA from formalin-fixed paraffin-embedded tumor tissues was extracted using the QIAamp DNA FFpE Tissue Kit (Qiagen) following manufacturer’s instructions. A nested, two-stage PCR approach to improve the sensitivity to detect methylated alleles was performed as previously described (Palmisano et al., 2000). Genomic DNA was subjected to bisulfite treatment using the Epitect Bisulfite Kit (Qiagen) and PCR was performed to amplify a 289 bp fragment of the MGMT promoter region. The primers recognize the bisulfite-modified template but do not discriminate between methylated and unmethylated alleles. The stage 1 PCR products were diluted 50-fold, and 5 μL was subjected to a stage 2 PCR in which primers specific to methylated or unmethylated template were used. Taq Gold polymerase (Thermo Fisher Scientific) in a 50 μL volume reaction was used in all PCRs. PCR amplification protocol for stage 1 was as follows: 95°C for 10 min, followed by denaturation at 95°C for 30 sec, annealing at 52°C for 30 sec and extension at 72°C for 30 sec for 40 cycles followed by a final extension at 72°C for 10 min. PCR amplification protocol for stage 2 was as follows: 95°C for 15 min, followed by denaturation at 95°C for 30 sec, annealing at 62°C for 30 sec and extension at 72°C for 30 sec for 2 cycles. Next, denaturation at 95°C for 30 sec, annealing at 60°C for 30 sec and extension at 72°C for 30 sec for 2 cycles was performed. Finally, denaturation at 95°C for 30 sec, annealing at 58°C for 30 sec and extension at 72°C for 30 sec for 36 cycles followed by a final extension at 72°C for 7 min was performed. Placental DNA treated with SssI methyltransferase (New England Biolabs) was used as a positive control for methylated alleles of MGMT, and DNA from normal lymphocytes was used as a negative control. Controls without DNA (blank) were used for each set of methylation-specific PCR assays. 7 μL of each methylation-specific PCR product was loaded directly into 3% agarose gel, stained with real safe (Durviz) and examined under ultraviolet illumination. Primers used to selectively amplify unmethylated or methylated MGMT gene in the stage 2 PCR were as previously described (Esteller et al., 2000; Hegi et al., 2005).

Primers (5’> 3’, forward; reverse):

- *MGMT* (stage 1): (GGATATGTTGGGATAGTT; CCAAAAACCCCAAACCC)
- *MGMT* unmethylated (stage 2): (TTTGTGTTTTGATGTTTGTAGGTTTTTGT; AACTCCACACTCTTCCAAAAACAAAACA)
- *MGMT* methylated (stage 2): (TTTCGACGTTCGTAGGTTTTCGC; GCACTCTTCCGAAAACGAAACG)

### *EGFR* mutational analysis

DNA was extracted from FFPE tissue samples and macrodissection was performed to ensure a content of at least 60% tumor cells. Samples were tested by real time PCR in a cobas z480 analyzer (Roche Diagnostics) using the cobas EGFR Mutation Test, which can detect mutations in exons 18, 19, 20 and 21 of the *EGFR* gene.

### qRT-PCR

Whole RNA was isolated using the RNAeasy Mini Kit (Qiagen) and was used (1,000 ng) to generate cDNA using iScript cDNA Synthesis Kit (1708891, Bio-Rad) according to manufacturer’s instructions. RNA obtained from mouse brains included microdissected established metastases from human BrM cells. Gene expression in the tumor was analyzed by using human primers using SYBR green gene expression assays (GoTaq qPCR Master Mix, A6002, Promega).

Primers (5’> 3’, forward; reverse):

- *HSP90AA1*: (AGATGACGACACATCACGCA; ACAGTGCACGTTACCCCAAT)
- *HSP90AB1*: (TGAGGAGGATGACAGCGGTA; TCAAAAAGGTCAAAGGGAGCC)
- *HSPA4* (HSP70): (GCAAGTGACTGCCATGCTTT; TAAGCAGAGTGGCCCATGTC)
- *HSPB2* (HSP27): (TAAACCTGGAAGCACCTCGG; ACATTGTGGACCATGCACCT)
- *HSF1*: (CCCTGATGCTGAACGACAGT; GGATAGGGGCCTCTCGTCTA)

### Transcriptomics of relapsed tumors

500ng of total RNA samples were used. Sample RNA Integrity numbers were 8.6 on average (range 5.9-9.5) when assayed on an Agilent 2100 Bioanalyzer. Sequencing libraries were prepared with the QuantSeq 3’ mRNA-Seq Library Prep Kit (FWD) for Illumina (Lexogen, Cat.No. 015) by following manufacturer’s instructions. Library generation is initiated by reverse transcription with oligodT priming, and a second strand synthesis is performed from random primers by a DNA polymerase. Primers from both steps contain Illumina-compatible sequences. Libraries are completed by PCR {This kit generates directional libraries stranded in the sense orientation: the read1, the only read in single read format, has the sense orientation (--library-type fr-secondstrand in TopHat, --stranded=yes in HTSeq)}. cDNA libraries are purified, applied to an Illumina flow cell for cluster generation and sequenced on an Illumina NextSeq 550 (with v2.5 reagent kits) by following manufacturer’s protocols. Eightyfive-base-pair single-end sequenced reads followed adapter and polyA tail removal as indicated by Lexogen. The resulting reads were fed to Xenome (Conway et al., 2012) to separate the xenograft-derived human and mouse reads. Human reads were analysed with the nextpresso (Graña et al., 2018) pipeline as follows: sequencing quality was checked with FastQC v0.11.0 (https://www.bioinformatics.babraham.ac.uk/projects/fastqc/). Reads were aligned to the human genome (GRCh38) with TopHat-2.0.10 (Trapnell et al., 2009) using Bowtie 1.0.0 (Langmead et al., 2009) and Samtools 0.1.19 (Li et al., 2009), allowing 3 mismatches and 20 multihits. The Gencode v29 gene annotation for GRCh38 was used. Read counts were obtained with HTSeq (Anders et al., 2015). Differential expression and normalization were performed with DESeq2 (Love et al., 2014), filtering out those genes where the normalized count value was lower than 2 in more than 50% of the samples. From the remaining genes, those that had an adjusted p-value below 0.05 FDR were selected. GSEAPreranked (Subramanian et al., 2005) was used to perform gene set enrichment analysis for several gene signatures on a pre-ranked gene list, setting 1000 gene set permutations. Only those gene sets with significant enrichment levels (FDR q-value < 0.25) were considered.

### Analysis of prognosis

A previously published RNA-sequencing dataset of 21 brain metastases from breast cancer patients with clinical annotation was downloaded (https://github.com/leeoesterreich/shiny-server/tree/master/apps/Paired_Mets) (Varešlija et al., 2019). *UBE4B* and *DDA1* gene expression was assessed using log_2_ transformed trimmed M of means (TMM)-normalized counts per million (log2(TMM-CPM +1)). Two groups of patients with low or high gene expression were delineated using the maximally selected rank statistics (Hothorn and Lausen, 2003), as implemented in the ‘survminer’ R package (Kassambara et al., 2019) and Kaplan-Meier survival curves were generated depicting survival post-brain metastasis.

### *In situ* proteomics

Fixed organotypic cultures were embedded in paraffin. 10 μm sections were placed on PET-membrane slides (415190-9051-000, Zeiss) pretreated with UV light. Slides were stained for 5 min in hematoxylin solution and 30 sec in eosin solution, and were left unmounted. Fully established brain metastases were isolated using the ArcturusXT™ Laser Capture Microdissection System (Thermo Scientific) and Arcturus^®^ CapSure^®^ Macro LCM Caps (Life Technologies) according to the manufacturer’s protocol. Each dissection was validated by inspection of the cap and the sample. At least 12 brain metastases per biological sample were dissected. Dissected samples were processed using the commercially available in-StageTip-NHS kit (PreOmics GmbH) according to the manufacturer’s protocol. Peptides were dissolved in HPLC-grade water containing 0.1% formic acid and 2% acetonitrile. Randomization for sample run order was applied and the samples were individually analyzed using shot-gun liquid chromatography tandem mass spectrometry (LC-MS/MS) on a high accuracy Orbitrap Fusion™ Lumos™ Tribrid™ Mass Spectrometer (Thermo Fisher) coupled to an Acquity M nanoflow system (Waters GmbH). Samples were analyzed using 120 minutes gradient, top12 loop count, mass range 350 to 1500 m/z and an Acquity UPLC^®^ M class 250 mm x 75 μM column. All raw files from LC-MS/MS were processed with MaxQuant (version 1.6.2.6) using the standard settings against a human protein database (UniProtKB/Swiss-Prot, 20,373 sequences) supplemented with contaminants. Label-free quantification was done with match between runs (match window of 0.7 min and alignment window of 20 min). Carbamidomethylation of cysteines was set as a fixed modification whereas oxidation of methionines and protein N-term acetylation as variable modifications. Minimal peptide length was set to 7 amino acids and a maximum of two tryptic missed-cleavages were allowed. Results were filtered at 0.01 FDR (peptide and protein level). Then, the “proteinGroups.txt” file was loaded in Prostar (v1.14) (Wieczorek et al., 2017) for further statistical analysis. Briefly, global normalization across samples was performed using the LOESS function and missing values were imputed using the algorithms slsa (for partially observed values) and detquantile (for values missing on an entire condition). Differential analysis was done using the empirical bayes statistics limma. Proteins with a p.value <0.05 and a log2 ratio >1 or <-1 were defined as deregulated. The FDR was estimated to be 14 % by Benjamini-Hochberg. Functional analysis was performed with the GSEApreranked function (biocarta, canonical pathways, GO, KEGG, OncogenicSignatures, Reactome, TFs) using the log2 ratios as the input file to identify top 25 upregulated and downregulated signatures defined by NES values, FDR<25% and P<0.01.

### Pharmacokinetics assay

Plasma and brain samples were collected 6h after oral administration of DEBIO-0932 (160 mg/kg) to brain metastases-bearing mice. Around 1 mL of blood was centrifuged at 3,000 r.p.m. for 10 min at 4 °C immediately after the extraction. Brain samples were homogenized in 4 volumes of H2O and sonicated for 10 min followed by centrifugation at 10,000 r.p.m. for 5 min. The supernatant was stored at −20 °C until processing. The extraction of DEBIO-0932 was achieved by solid-phase extraction followed by high-performance liquid chromatography/ tandem mass spectrometry (Agilent 1100, Sciex QTRAP 5500 System) analysis. The amount of DEBIO-0932 in each sample was quantified based on calibration curves generated using standards of known concentrations of DEBIO-0932. For the conversion of brain concentrations in ng/g to ng/mL, a tissue density of 1 was assumed.

### Magnetic resonance imaging

MRI studies were carried out in a Bruker Biospec 70/20 scanner using a combination of a linear coil (for transmission) with a mouse head phase array coil (for reception). Animals were anesthetized with sevoflurane (5% for induction and 3% for maintenance) and placed in an MRI-adapted stereotaxic holder with a water circulating blanket to maintain body temperature. Respiration and body temperature were continuously monitored. As anatomical reference, a T2-weighted sequence was acquired (TR= 4600 ms; TE, 65 ms; α = 90°; FOV = 1.5 x 1.5 cm; matrix = 192×192; slice thickness = 0.5 mm, number of slices = 30). Then, a T1 sequence was acquired (TR= 472.610 ms; TE, 3.648 ms; α = 30°; FOV = 1.5 x 1.5 cm; matrix = 192×192; slice thickness = 0.5 mm, number of slices = 30) before and after intravenous administration of 200 μL of Gadovist (1mmol/ml, Bayer AG).

### Image acquisition and analysis

Immunofluorescence images were acquired with a Leica SP5 up-right confocal microscope x5, x10, x20, x40 and x63 objectives and analyzed with ImageJ software and Definiens developer XD 2.5. Immunohistochemistry images were captured with the Zen Blue Software v3.1 (Zeiss) and whole slides were acquired with a slide scanner (AxioScan Z1, Zeiss). For histological quantification of brain metastases at endpoint (5 weeks after intracardiac inoculation of cancer cells), only lesions showing solid and compact distribution of cancer cells were considered as established metastases.

### Statistical analysis

Data are represented as the mean ± s.e.m. Comparisons between two experimental groups were analyzed with unpaired, two-tailed Student’s t-test. Survival analysis was done with log-rank (Mantel-Cox) test.

## Supplemental figure legends

**Figure S1.**
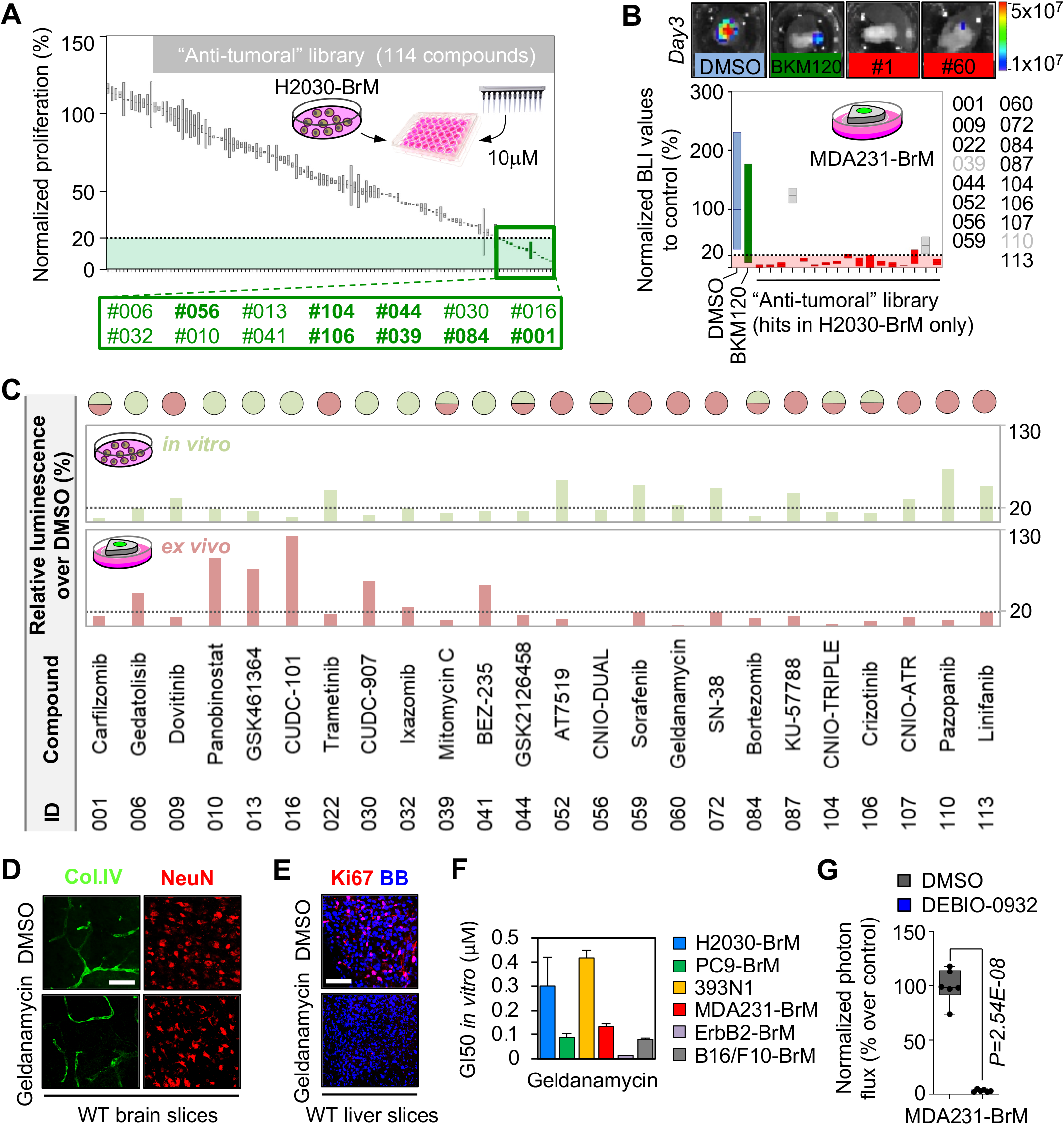
A chemical library applied to METPlatform identifies potential vulnerabilities of brain metastasis. **(A)** Quantification of the proliferation of H2030-BrM cells at day 3 normalized to the cells treated with DMSO measured with CellTiter-Glo^®^. Green: hits, compounds with ≤20% proliferation; gray: compounds with >20% proliferation. Values are shown in box-and-whisker plots where the line in the box corresponds to the mean. Each experimental compound of the library was assayed by duplicate. Hits highlighted in bold were common to the *ex vivo* screening (Fig. 1 B). **(B)** Quantification of the bioluminescence signal from MDA231-BrM established brain metastases in organotypic culture after 3 days in culture. Values were normalized by the level of bioluminescence at day 0 for each culture (before the addition of DMSO or any compound). Final data is shown in percentage respect to reference, the organotypic cultures treated with DMSO. Blue: DMSO-treated organotypic cultures; red: hits, compounds with normalized BLI ≤20%; green: BKM120; gray: compounds with normalized BLI >20%. Values are shown in box-and-whisker plots where the line in the box corresponds to the mean. Whiskers go from the minimum to the maximum value (n=14 DMSO; n=13 BKM120-treated organotypic cultures; each experimental compound was assayed by duplicate, 4 independent experiments). **(C)** Detailed representation of the data shown in Fig.1 B and Fig. S1 A indicating relative proliferation values (%) of H2030-BrM cells *ex vivo* (established brain metastases, light red) and *in vitro* (green) treated with compounds of the anti-tumoral library (compounds were assayed by duplicate in each assay). All hits for any condition are shown. The circles of the top indicate whether a given compound was effective (<20% luminescence respect to control) *ex vivo* (light red circle), *in vitro* (green circle) or both (light red and green circles). **(D)** Representative wild type brain slices treated with DMSO or the HSP90 inhibitor geldanamycin stained with anti-Col.IV (endothelial cells) and anti-NeuN (neurons). Scale bar, 50 μm. **(E)** Representative wild type liver slices treated with DMSO or the HSP90 inhibitor geldanamycin and stained with anti-Ki67 to score proliferation. BB: bisbenzamide. Scale bar, 50 μm. **(F)** Quantification of GI50 values of geldanamycin in a panel of BrM cell lines *in vitro* from various primary origins and oncogenomic profiles. Serial concentrations of geldanamycin were assayed by duplicate and GI50 was calculated from a viability curve normalized to DMSO treated cells of the corresponding cell line. Values are shown as mean + s.e.m. (each concentration was assayed by duplicate for each cell line). **(G)** Quantification of the bioluminescence signal emitted by MDA231-BrM established metastases in organotypic cultures incubated in the presence of DEBIO-0932 (1μM) during 3 days. Bioluminescence at day 3 is normalized by the initial value obtained at day 0 and quantified relative to the organotypic cultures treated with DMSO. Day 0 is considered right before addition of the treatment or DMSO. Values are shown in box-and-whisker plots where each dot is an organotypic culture and the line in the box corresponds to the median. Whiskers go from the minimum to the maximum value (n=6 organotypic cultures per experimental condition, 1 independent experiment). *P* value was calculated using two-tailed t-test.

**Figure S3.**
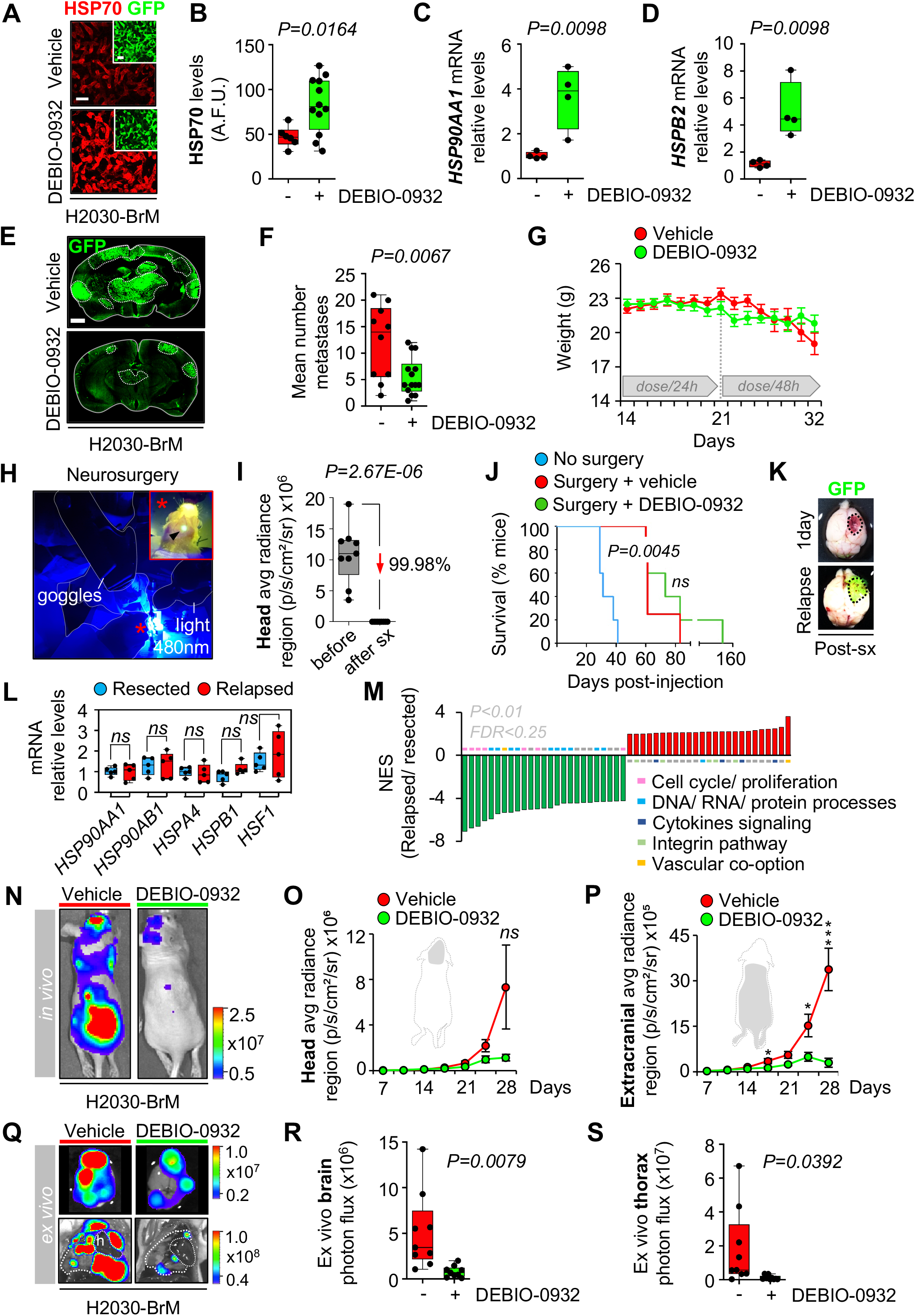
Inhibition of HSP90 impairs clinically-relevant stages of brain metastasis. **(A)** Representative images showing HSP70 levels in brain metastases (generated by intracardiac inoculation of H2030-BrM) found at endpoint of vehicle and DEBIO-0932 treated animals. Scale bars, 75 μm. **(B)** Quantification of HSP70 levels shown in (A) in arbitrary fluorescent units (A.F.U.). Values are shown in box-and-whisker plots where each dot is a metastatic lesion and the line in the box corresponds to the median. Whiskers go from the minimum to the maximum value (n=6-12 metastatic lesions from 3-6 brains per condition). P value was calculated using two-tailed t-test. **(C-D)** *HSP90AA1* **(C)** and *HSPB2* **(D)** expression levels obtained by qRT-PCR of H2030-BrM brain metastases obtained at endpoint of vehicle and DEBIO-0932 treated animals. Values are shown in box-and-whisker plots where every dot represents a different animal and the line in the box corresponds to the median. Whiskers go from the minimum to the maximum value (n=4 mice per experimental condition). *P* value was calculated using two-tailed t-test. **(E)** Representative sections of brains from vehicle and DEBIO-0932 treated mice 5 weeks (experimental endpoint) after intracardiac inoculation of H2030-BrM cells. DEBIO-0932 treatment started 2 weeks after inoculation of cancer cells. The dotted lines surround the metastases (GFP+). Scale bar, 1mm. **(F)** Quantification of established metastases found in vehicle and DEBIO-0932 treated brains from panel (A). Values are shown in box-and-whisker plots where every dot represents a different brain and the line in the box corresponds to the median. Whiskers go from the minimum to the maximum value (vehicle: n=10 brains; DEBIO-0932: n=14 brains). *P* value was calculated using two-tailed t-test. **(G)** Animal weight from vehicle and DEBIO-0932 treated mice during the treatment period. DEBIO-0932 treatment started 2 weeks (day 14) after inoculation of cancer cells and was maintained for 3 weeks, once every 24h during the first week and once every 48h during the two following weeks. Values are shown as mean ± s.e.m. (n=9 vehicle and n=10 DEBIO-0932 treated mice). **(H)** Detailed image of the neurosurgery procedure that visualizes the GFP+ brain tumor (high magnification) with a 480nm light source and goggles equipped with emission filters. **(I)** Quantification of BLI values before and one day after neurosurgery. Values are shown in box-and-whisker plots where every dot represents a different animal and the line in the box corresponds to the median. Whiskers go from the minimum to the maximum value (n=9 mice before and after surgery). *P* value was calculated using two-tailed t-test. **(J)** Kaplan-Meier curve showing survival proportions of mice without (blue line, n=5) and with surgery and vehicle (red line, n=4) or DEBIO-0932 (green line, n=5). *P* value was calculated using log-rank (Mantel-Cox) test. **(K)** Representative images of brains one day after neurosurgery and at the endpoint when relapsed tumor is fully developed and GFP+ cancer cells could be easily detected. **(L)** qRT-PCR of H2030-BrM brain metastases obtained from animals during neurosurgery compared to relapsed metastases from the corresponding animals. A panel of five genes related to HSP90 pathway is evaluated. Values are shown in box-and-whisker plots where every dot represents a different animal and the line in the box corresponds to the median. Whiskers go from the minimum to the maximum value (n=5 mice per experimental condition). *P* value was calculated using two-tailed t-test. **(M)** GSEA of top 25 up-(red) and downregulated (green) signatures comparing matched relapsed and resected brain metastases from animals receiving neurosurgery. **(N)** Representative images of mice treated with DEBIO-0932 (160 mg/kg, o.g.) starting at 7 days after intracardiac inoculation of H2030-BrM cells. Treatment was maintained until the endpoint of the experiment at 4 weeks. **(O-P)** Quantification of metastatic progression as measured by *in vivo* BLI of head (O) and extracranial region (P) of animals. Values are shown as mean ± s.e.m. (n=9 vehicle and n=9 DEBIO-0932 treated mice, 2 independent experiments). *P* value was calculated using two-tailed t-test (*P* values: \**P*<0.05, \*\*\**P*<0.001). **(Q)** Representative images of brains and thorax from vehicle and DEBIO-0932 treated mice at the endpoint of the experiment. (R-S) Quantification of *ex vivo* BLI of brains (R) and thoracic regions (S) at the endpoint of the experiment. Values are shown in box-and-whisker plots where every dot represents a different animal and the line in the box corresponds to the median. Whiskers go from the minimum to the maximum value (n=9 vehicle and n=9 DEBIO-0932 treated mice, 2 independent experiments). *P* value was calculated using two-tailed t-test.

## References

Anders, S., Pyl, P.T., and Huber, W. (2015). HTSeq — a Python framework to work with high-throughput sequencing data. Bioinformatics 31, 166–169.

Antonova, A., Hummel, B., Khavaran, A., Redhaber, D.M., Aprile-Garcia, F., Rawat, P., Gundel, K., Schneck, M., Hansen, E.C., Mitschke, J., et al. (2019). Heat-Shock Protein 90 Controls the Expression of Cell-Cycle Genes by Stabilizing Metazoan-Specific Host-Cell Factor HCFC1. Cell Rep. 29, 1645–1659.e9.

Arvold, N.D., Lee, E.Q., Mehta, M.P., Margolin, K., Alexander, B.M., Lin, N.U., Anders, C.K., Soffietti, R., Camidge, D.R., Vogelbaum, M.A., et al. (2016). Updates in the management of brain metastases. Neuro. Oncol. 18, 1043–1065.

Bao, R., Lai, C.-J., Qu, H., Wang, D., Yin, L., Zifcak, B., Atoyan, R., Wang, J., Samson, M., Forrester, J., et al. (2009). CUDC-305, a novel synthetic HSP90 inhibitor with unique pharmacologic properties for cancer therapy. Clin. Cancer Res. 15, 4046–4057.

Basnet, H., Tian, L., Ganesh, K., Huang, Y.-H., Macalinao, D.G., Brogi, E., Finley, L.W., and Massagué, J. (2019). Flura-seq identifies organ-specific metabolic adaptations during early metastatic colonization. Elife 8.

Bejarano, L., Bosso, G., Louzame, J., Serrano, R., Gómez-Casero, E., Martínez-Torrecuadrada, J., Martínez, S., Blanco-Aparicio, C., Pastor, J., and Blasco, M.A. (2019). Multiple cancer pathways regulate telomere protection. EMBO Mol. Med. 11, e10292.

Boire, A., Brastianos, P.K., Garzia, L., and Valiente, M. (2020). Brain metastasis. Nat. Rev. Cancer 20, 4–11.

Bos, P.D., Zhang, X.H.-F., Nadal, C., Shu, W., Gomis, R.R., Nguyen, D.X., Minn, A.J., van de Vijver, M.J., Gerald, W.L., Foekens, J.A., et al. (2009). Genes that mediate breast cancer metastasis to the brain. Nature 459, 1005–1009.

Brastianos, P.K., Carter, S.L., Santagata, S., Cahill, D.P., Taylor-Weiner, A., Jones, R.T., Van Allen, E.M., Lawrence, M.S., Horowitz, P.M., Cibulskis, K., et al. (2015). Genomic characterization of brain metastases reveals branched evolution and potential therapeutic targets. Cancer Discov. 5, 1164–1177.

Butler, M., Pongor, L., Su, Y.-T., Xi, L., Raffeld, M., Quezado, M., Trepel, J., Aldape, K., Pommier, Y., and Wu, J. (2020). MGMT status as a clinical biomarker in glioblastoma. Trends Cancer 6, 380–391.

Camidge, D.R., Lee, E.Q., Lin, N.U., Margolin, K., Ahluwalia, M.S., Bendszus, M., Chang, S.M., Dancey, J., de Vries, E.G.E., Harris, G.J., et al. (2018). Clinical trial design for systemic agents in patients with brain metastases from solid tumours: a guideline by the Response Assessment in Neuro-Oncology Brain Metastases working group. Lancet Oncol. 19, e20–e32.

Cedrés, S., Felip, E., Cruz, C., Martinez de Castro, A., Pardo, N., Navarro, A., Martinez-Marti, A., Remon, J., Zeron-Medina, J., Balmaña, J., et al. (2018). Activity of HSP90 inhibiton in a metastatic lung cancer patient with a germline BRCA1 mutation. J. Natl. Cancer Inst. 110, 914–917.

Chen, P.-H., Chang, J.T., Li, L.-A., Tsai, H.-T., Shen, M.-Y., and Lin, P. (2013). Aryl hydrocarbon receptor is a target of 17-Allylamino-17-demethoxygeldanamycin and enhances its anticancer activity in lung adenocarcinoma cells. Mol. Pharmacol. 83, 605–612.

Chen, Q., Boire, A., Jin, X., Valiente, M., Er, E.E., Lopez-Soto, A., Jacob, L., Patwa, R., Shah, H., Xu, K., et al. (2016). Carcinoma-astrocyte gap junctions promote brain metastasis by CGAMP transfer. Nature 533, 493–498.

Cheng, L., Yang, Q., Li, C., Dai, L., Yang, Y., Wang, Q., Ding, Y., Zhang, J., Liu, L., Zhang, S., et al. (2017). DDA1, a novel oncogene, promotes lung cancer progression through regulation of cell cycle. J. Cell Mol. Med. 21, 1532–1544.

Conway, T., Wazny, J., Bromage, A., Tymms, M., Sooraj, D., Williams, E.D., and Beresford-Smith, B. (2012). Xenome--a tool for classifying reads from xenograft samples. Bioinformatics 28, i172–8.

DeVorkin, L., and Lum, J.J. (2014). Strategies to block autophagy in tumor cells. In Autophagy: cancer, other pathologies, inflammation, immunity, infection, and aging, (Elsevier), pp. 121–130.

Dimas, D.T., Perlepe, C.D., Sergentanis, T.N., Misitzis, I., Kontzoglou, K., Patsouris, E., Kouraklis, G., Psaltopoulou, T., and Nonni, A. (2018). The Prognostic Significance of Hsp70/Hsp90 Expression in Breast Cancer: A Systematic Review and Meta-analysis. Anticancer Res. 38, 1551–1562.

Du, C., Wu, H., and Leng, R.P. (2016). UBE4B targets phosphorylated p53 at serines 15 and 392 for degradation. Oncotarget 7, 2823–2836.

Echeverría, P.C., Bernthaler, A., Dupuis, P., Mayer, B., and Picard, D. (2011). An interaction network predicted from public data as a discovery tool: application to the Hsp90 molecular chaperone machine. PLoS One 6, e26044.

Er, E.E., Valiente, M., Ganesh, K., Zou, Y., Agrawal, S., Hu, J., Griscom, B., Rosenblum, M., Boire, A., Brogi, E., et al. (2018). Pericyte-like spreading by disseminated cancer cells activates YAP and MRTF for metastatic colonization. Nat. Cell Biol. 20, 966–978.

Esteller, M., Garcia-Foncillas, J., Andion, E., Goodman, S.N., Hidalgo, O.F., Vanaclocha, V., Baylin, S.B., and Herman, J.G. (2000). Inactivation of the DNA-repair gene MGMT and the clinical response of gliomas to alkylating agents. N. Engl. J. Med. 343, 1350–1354.

Fionda, C., Soriani, A., Malgarini, G., Iannitto, M.L., Santoni, A., and Cippitelli, M. (2009). Heat shock protein-90 inhibitors increase MHC class I-related chain A and B ligand expression on multiple myeloma cells and their ability to trigger NK cell degranulation. J. Immunol. 183, 4385–4394.

Gallegos Ruiz, M.I., Floor, K., Roepman, P., Rodriguez, J.A., Meijer, G.A., Mooi, W.J., Jassem, E., Niklinski, J., Muley, T., van Zandwijk, N., et al. (2008). Integration of gene dosage and gene expression in non-small cell lung cancer, identification of HSP90 as potential target. PLoS One 3, e0001722.

Gao, H., Korn, J.M., Ferretti, S., Monahan, J.E., Wang, Y., Singh, M., Zhang, C., Schnell, C., Yang, G., Zhang, Y., et al. (2015). High-throughput screening using patient-derived tumor xenografts to predict clinical trial drug response. Nat. Med. 21, 1318–1325.

García-Gómez, P., and Valiente, M. (2020). Vascular co-option in brain metastasis. Angiogenesis 23, 3–8.

Garralda, E., Paz, K., López-Casas, P.P., Jones, S., Katz, A., Kann, L.M., López-Rios, F., Sarno, F., Al-Shahrour, F., Vasquez, D., et al. (2014). Integrated next-generation sequencing and avatar mouse models for personalized cancer treatment. Clin. Cancer Res. 20, 2476–2484.

Graña, O., Rubio-Camarillo, M., Fdez-Riverola, F., Pisano, D.G., and Glez-Peña, D. (2018). Nextpresso: next generation sequencing expression analysis pipeline. Curr. Bioinform. 13, 583–591.

Gril, B., Palmieri, D., Qian, Y., Anwar, T., Liewehr, D.J., Steinberg, S.M., Andreu, Z., Masana, D., Fernández, P., Steeg, P.S., et al. (2013). Pazopanib inhibits the activation of PDGFRβ-expressing astrocytes in the brain metastatic microenvironment of breast cancer cells. Am. J. Pathol. 182, 2368–2379.

He, W., Ye, X., Huang, X., Lel, W., You, L., Wang, L., Chen, X., and Qian, W. (2016). Hsp90 inhibitor, BIIB021, induces apoptosis and autophagy by regulating mTOR-Ulk1 pathway in imatinib-sensitive and - resistant chronic myeloid leukemia cells. Int. J. Oncol. 48, 1710–1720.

Hegi, M.E., Diserens, A.-C., Gorlia, T., Hamou, M.-F., de Tribolet, N., Weller, M., Kros, J.M., Hainfellner, J.A., Mason, W., Mariani, L., et al. (2005). MGMT gene silencing and benefit from temozolomide in glioblastoma. N. Engl. J. Med. 352, 997–1003.

Hirata, E., and Sahai, E. (2017). Tumor microenvironment and differential responses to therapy. Cold Spring Harb. Perspect. Med. 7.

Hothorn, T., and Lausen, B. (2003). On the exact distribution of maximally selected rank statistics. Comput. Stat. Data Anal. 43, 121–137.

Humpel, C. (2015). Organotypic brain slice cultures: A review. Neuroscience 305, 86–98.

Isambert, N., Delord, J.P., Soria, J.C., Hollebecque, A., Gomez-Roca, C., Purcea, D., Rouits, E., Belli, R., and Fumoleau, P. (2015). Debio0932, a second-generation oral heat shock protein (HSP) inhibitor, in patients with advanced cancer-results of a first-in-man dose-escalation study with a fixed-dose extension phase. Ann. Oncol. 26, 1005–1011.

Kaizuka, T., Morishita, H., Hama, Y., Tsukamoto, S., Matsui, T., Toyota, Y., Kodama, A., Ishihara, T., Mizushima, T., and Mizushima, N. (2016). An Autophagic Flux Probe that Releases an Internal Control. Mol. Cell 64, 835–849.

Kassambara, A., Kosinski, M., Biecek, P., and Fabian, S. (2019). survminer: Drawing Survival Curves using “ggplot2”. 2019. R package version 0.4 4.

Kawabe, M., Mandic, M., Taylor, J.L., Vasquez, C.A., Wesa, A.K., Neckers, L.M., and Storkus, W.J. (2009). Heat shock protein 90 inhibitor 17-dimethylaminoethylamino-17-demethoxygeldanamycin enhances EphA2+ tumor cell recognition by specific CD8+ T cells. Cancer Res. 69, 6995–7003.

Keeling, D.J., Herslöf, M., Ryberg, B., Sjögren, S., and Sölvell, L. (1997). Vacuolar H(+)-ATPases. Targets for drug discovery? Ann. N. Y. Acad. Sci. 834, 600–608.

Kim, K., Lee, H.W., Lee, E.H., Park, M.-I., Lee, J.S., Kim, M.-S., Kim, K., Roh, M.S., Pak, M.G., Oh, J.E., et al. (2019). Differential expression of HSP90 isoforms and their correlations with clinicopathologic factors in patients with colorectal cancer. Int J Clin Exp Pathol 12, 978–986.

Langmead, B., Trapnell, C., Pop, M., and Salzberg, S.L. (2009). Ultrafast and memory-efficient alignment of short DNA sequences to the human genome. Genome Biol. 10, R25.

Lee, J.-K., Liu, Z., Sa, J.K., Shin, S., Wang, J., Bordyuh, M., Cho, H.J., Elliott, O., Chu, T., Choi, S.W., et al. (2018). Pharmacogenomic landscape of patient-derived tumor cells informs precision oncology therapy. Nat. Genet. 50, 1399–1411.

Levy, J.M.M., Towers, C.G., and Thorburn, A. (2017). Targeting autophagy in cancer. Nat. Rev. Cancer 17, 528–542.

Li, H., Handsaker, B., Wysoker, A., Fennell, T., Ruan, J., Homer, N., Marth, G., Abecasis, G., Durbin, R., and 1000 Genome Project Data Processing Subgroup (2009). The Sequence Alignment/Map format and SAMtools. Bioinformatics 25, 2078–2079.

Lin, N.U., Lee, E.Q., Aoyama, H., Barani, I.J., Baumert, B.G., Brown, P.D., Camidge, D.R., Chang, S.M., Dancey, J., Gaspar, L.E., et al. (2013a). Challenges relating to solid tumour brain metastases in clinical trials, part 1: patient population, response, and progression. A report from the RANO group. Lancet Oncol. 14, e396–406.

Lin, N.U., Wefel, J.S., Lee, E.Q., Schiff, D., van den Bent, M.J., Soffietti, R., Suh, J.H., Vogelbaum, M.A., Mehta, M.P., Dancey, J., et al. (2013b). Challenges relating to solid tumour brain metastases in clinical trials, part 2: neurocognitive, neurological, and quality-of-life outcomes. A report from the RANO group. Lancet Oncol. 14, e407–16.

Lin, N.U., Lee, E.Q., Aoyama, H., Barani, I.J., Barboriak, D.P., Baumert, B.G., Bendszus, M., Brown, P.D., Camidge, D.R., Chang, S.M., et al. (2015). Response assessment criteria for brain metastases: proposal from the RANO group. Lancet Oncol. 16, e270–8.

Liu, K.-S., Liu, H., Qi, J.-H., Liu, Q.-Y., Liu, Z., Xia, M., Xing, G.-W., Wang, S.-X., and Wang, Y.-F. (2012). SNX-2112, an Hsp90 inhibitor, induces apoptosis and autophagy via degradation of Hsp90 client proteins in human melanoma A-375 cells. Cancer Lett. 318, 180–188.

Love, M.I., Huber, W., and Anders, S. (2014). Moderated estimation of fold change and” dispersion for RNA-seq data with DESeq2. Genome Biol. 15, 550.

Mauvezin, C., Nagy, P., Juhász, G., and Neufeld, T.P. (2015). Autophagosome-lysosome fusion is independent of V-ATPase-mediated acidification. Nat. Commun. 6, 7007.

McCarthy, M.M., Pick, E., Kluger, Y., Gould-Rothberg, B., Lazova, R., Camp, R.L., Rimm, D.L., and Kluger, H.M. (2008). HSP90 as a marker of progression in melanoma. Ann. Oncol. 19, 590–594.

Moravan, M.J., Fecci, P.E., Anders, C.K., Clarke, J.M., Salama, A.K.S., Adamson, J.D., Floyd, S.R., Torok, J.A., Salama, J.K., Sampson, J.H., et al. (2020). Current multidisciplinary management of brain metastases. Cancer.

Mori, M., Hitora, T., Nakamura, O., Yamagami, Y., Horie, R., Nishimura, H., and Yamamoto, T. (2015). Hsp90 inhibitor induces autophagy and apoptosis in osteosarcoma cells. Int. J. Oncol. 46, 47–54.

Morrissy, A.S., Garzia, L., Shih, D.J.H., Zuyderduyn, S., Huang, X., Skowron, P., Remke, M., Cavalli, F.M.G., Ramaswamy, V., Lindsay, P.E., et al. (2016). Divergent clonal selection dominates medulloblastoma at recurrence. Nature 529, 351–357.

Murray, I.A., Patterson, A.D., and Perdew, G.H. (2014). Aryl hydrocarbon receptor ligands in cancer: friend and foe. Nat. Rev. Cancer 14, 801–814.

Nadanaciva, S., Lu, S., Gebhard, D.F., Jessen, B.A., Pennie, W.D., and Will, Y. (2011). A high content screening assay for identifying lysosomotropic compounds. Toxicol. In Vitro 25, 715–723.

Nahed, B.V., Alvarez-Breckenridge, C., Brastianos, P.K., Shih, H., Sloan, A., Ammirati, M., Kuo, J.S., Ryken, T.C., Kalkanis, S.N., and Olson, J.J. (2019). Congress of Neurological Surgeons Systematic Review and Evidence-Based Guidelines on the Role of Surgery in the Management of Adults With Metastatic Brain Tumors. Neurosurgery 84, E152–E155.

Nanni, P., Nicoletti, G., Palladini, A., Croci, S., Murgo, A., Ianzano, M.L., Grosso, V., Stivani, V., Antognoli, A., Lamolinara, A., et al. (2012). Multiorgan metastasis of human HER-2+ breast cancer in Rag2-/-;Il2rg-/-mice and treatment with PI3K inhibitor. PLoS One 7, e39626.

Neckers, L., and Workman, P. (2012). Hsp90 molecular chaperone inhibitors: are we there yet? Clin. Cancer Res. 18, 64–76.

Nguyen, D.X., Chiang, A.C., Zhang, X.H.-F., Kim, J.Y., Kris, M.G., Ladanyi, M., Gerald, W.L., and Massagué, J. (2009). WNT/TCF signaling through LEF1 and HOXB9 mediates lung adenocarcinoma metastasis. Cell 138, 51–62.

Orgaz, J.L., Crosas-Molist, E., Sadok, A., Perdrix-Rosell, A., Maiques, O., Rodriguez-Hernandez, I., Monger, J., Mele, S., Georgouli, M., Bridgeman, V., et al. (2020). Myosin II reactivation and cytoskeletal remodeling as a hallmark and a vulnerability in melanoma therapy resistance. Cancer Cell 37, 85–103.e9.

Osswald, M., Blaes, J., Liao, Y., Solecki, G., Gömmel, M., Berghoff, A.S., Salphati, L., Wallin, J.J., Phillips, H.S., Wick, W., et al. (2016). Impact of Blood-Brain Barrier Integrity on Tumor Growth and Therapy Response in Brain Metastases. Clin. Cancer Res. 22, 6078–6087.

Palmisano, W.A., Divine, K.K., Saccomanno, G., Gilliland, F.D., Baylin, S.B., Herman, J.G., and Belinsky, S.A. (2000). Predicting lung cancer by detecting aberrant promoter methylation in sputum. Cancer Res. 60, 5954–5958.

Park, E.S., Kim, S.J., Kim, S.W., Yoon, S.-L., Leem, S.-H., Kim, S.-B., Kim, S.M., Park, Y.-Y., Cheong, J.-H., Woo, H.G., et al. (2011). Cross-species hybridization of microarrays for studying tumor transcriptome of brain metastasis. Proc. Natl. Acad. Sci. USA 108, 17456–17461.

Pick, E., Kluger, Y., Giltnane, J.M., Moeder, C., Camp, R.L., Rimm, D.L., and Kluger, H.M. (2007). High HSP90 expression is associated with decreased survival in breast cancer. Cancer Res. 67, 2932–2937.

Pistilli, B., Pluard, T., Urruticoechea, A., Farci, D., Kong, A., Bachelot, T., Chan, S., Han, H.S., Jerusalem, G., Urban, P., et al. (2018). Phase II study of buparlisib (BKM120) and trastuzumab in patients with HER2+ locally advanced or metastatic breast cancer resistant to trastuzumab-based therapy. Breast Cancer Res. Treat. 168, 357–364.

Priego, N., Zhu, L., Monteiro, C., Mulders, M., Wasilewski, D., Bindeman, W., Doglio, L., Martínez, L., Martínez-Saez, E., Ramón Y Cajal, S., et al. (2018). STAT3 labels a subpopulation of reactive astrocytes required for brain metastasis. Nat. Med. 24, 1024–1035.

Samarasinghe, B., Wales, C.T.K., Taylor, F.R., and Jacobs, A.T. (2014). Heat shock factor 1 confers resistance to Hsp90 inhibitors through p62/SQSTM1 expression and promotion of autophagic flux. Biochem. Pharmacol. 87, 445–455.

Saxena, D., Sharma, A., Siddiqui, M.H., and Kumar, R. (2019). Blood brain barrier permeability prediction using machine learning techniques: an update. Curr. Pharm. Biotechnol. 20, 1163–1171.

Schopf, F.H., Biebl, M.M., and Buchner, J. (2017). The HSP90 chaperone machinery. Nat. Rev. Mol. Cell Biol. 18, 345–360.

Sevenich, L., Bowman, R.L., Mason, S.D., Quail, D.F., Rapaport, F., Elie, B.T., Brogi, E., Brastianos, P.K., Hahn, W.C., Holsinger, L.J., et al. (2014). Analysis of tumour- and stroma-supplied proteolytic networks reveals a brainmetastasis-promoting role for cathepsin S. Nat. Cell Biol. 16, 876–888.

Shamir, E.R., and Ewald, A.J. (2014). Three-dimensional organotypic culture: experimental models of mammalian biology and disease. Nat. Rev. Mol. Cell Biol. 15, 647–664.

Stupp, R., Mason, W.P., van den Bent, M.J., Weller, M., Fisher, B., Taphoorn, M.J.B., Belanger, K., Brandes, A.A., Marosi, C., Bogdahn, U., et al. (2005). Radiotherapy plus concomitant and adjuvant temozolomide for glioblastoma. N. Engl. J. Med. 352, 987–996.

Stupp, R., Hegi, M.E., Mason, W.P., van den Bent, M.J., Taphoorn, M.J.B., Janzer, R.C., Ludwin, S.K., Allgeier, A., Fisher, B., Belanger, K., et al. (2009). Effects of radiotherapy with concomitant and adjuvant temozolomide versus radiotherapy alone on survival in glioblastoma in a randomised phase III study: 5-year analysis of the EORTC-NCIC trial. Lancet Oncol. 10, 459–466.

Su, J.-M., Hsu, Y.-Y., Lin, P., and Chang, H. (2016). Nuclear Accumulation of Heat-shock Protein 90 Is Associated with Poor Survival and Metastasis in Patients with Non-small Cell Lung Cancer. Anticancer Res. 36, 2197–2203.

Subramanian, A., Tamayo, P., Mootha, V.K., Mukherjee, S., Ebert, B.L., Gillette, M.A., Paulovich, A., Pomeroy, S.L., Golub, T.R., Lander, E.S., et al. (2005). Gene set enrichment analysis: a knowledge-based approach for interpreting genome-wide expression profiles. Proc. Natl. Acad. Sci. USA 102, 15545–15550.

Suh, J.H., Kotecha, R., Chao, S.T., Ahluwalia, M.S., Sahgal, A., and Chang, E.L. (2020). Current approaches to the management of brain metastases. Nat. Rev. Clin. Oncol. 17, 279–299.

Sui, X., Chen, R., Wang, Z., Huang, Z., Kong, N., Zhang, M., Han, W., Lou, F., Yang, J., Zhang, Q., et al. (2013). Autophagy and chemotherapy resistance: a promising therapeutic target for cancer treatment. Cell Death Dis. 4, e838.

Supko, J.G., Hickman, R.L., Grever, M.R., and Malspeis, L. (1995). Preclinical pharmacologic evaluation of geldanamycin as an antitumor agent. Cancer Chemother. Pharmacol. 36, 305–315.

Trapnell, C., Pachter, L., and Salzberg, S.L. (2009). TopHat: discovering splice junctions with RNA-Seq. Bioinformatics 25, 1105–1111.

Trepel, J., Mollapour, M., Giaccone, G., and Neckers, L. (2010). Targeting the dynamic HSP90 complex in cancer. Nat. Rev. Cancer 10, 537–549.

Tsimberidou, A.M., Letourneau, K., Wen, S., Wheler, J., Hong, D., Naing, A., Iskander, N.G., Uehara, C., and Kurzrock, R. (2011). Phase I clinical trial outcomes in 93 patients with brain metastases: the MD anderson cancer center experience. Clin. Cancer Res. 17, 4110–4118.

Valiente, M. (2020). Brain Metastasis Cell Lines Panel: a public resource of organotropic cell lines. Cancer Res.

Valiente, M., Obenauf, A.C., Jin, X., Chen, Q., Zhang, X.H.-F., Lee, D.J., Chaft, J.E., Kris, M.G., Huse, J.T., Brogi, E., et al. (2014). Serpins promote cancer cell survival and vascular co-option in brain metastasis. Cell 156, 1002–1016.

Valiente, M., Ahluwalia, M.S., Boire, A., Brastianos, P.K., Goldberg, S.B., Lee, E.Q., Le Rhun, E., Preusser, M., Winkler, F., and Soffietti, R. (2018). The evolving landscape of brain metastasis. Trends Cancer 4, 176–196.

Valiente, M., Van Swearingen, A.E.D., Anders, C.K., Bairoch, A., Boire, A., Bos, P.D., Cittelly, D.M., Erez, N., Ferraro, G.B., Fukumura, D., et al. (2020). Brain Metastasis Cell Lines Panel: a public resource of organotropic cell lines. Cancer Res.

Varešlija, D., Priedigkeit, N., Fagan, A., Purcell, S., Cosgrove, N., O’Halloran, P.J., Ward, E., Cocchiglia, S., Hartmaier, R., Castro, C.A., et al. (2019). Transcriptome characterization of matched primary breast and brain metastatic tumors to detect novel actionable targets. J. Natl. Cancer Inst. 111, 388–398.

Vlachogiannis, G., Hedayat, S., Vatsiou, A., Jamin, Y., Fernández-Mateos, J., Khan, K., Lampis, A., Eason, K., Huntingford, I., Burke, R., et al. (2018). Patient-derived organoids model treatment response of metastatic gastrointestinal cancers. Science 359, 920–926.

Whitesell, L., and Lindquist, S.L. (2005). HSP90 and the chaperoning of cancer. Nat. Rev. Cancer 5, 761–772.

Wieczorek, S., Combes, F., Lazar, C., Giai Gianetto, Q., Gatto, L., Dorffer, A., Hesse, A.-M., Couté, Y., Ferro, M., Bruley, C., et al. (2017). DAPAR & ProStaR: software to perform statistical analyses in quantitative discovery proteomics. Bioinformatics 33, 135–136.

Zhao, Y., Li, K., Zhao, B., and Su, L. (2019). HSP90 inhibitor DPB induces autophagy and more effectively apoptosis in A549 cells combined with autophagy inhibitors. In Vitro Cell Dev Biol Anim 55, 349–354.

Zhu, L., and Valiente, M. Organotypic brain cultures for metastasis research. Neuromethods (In press).

